# Identification and validation of drugs for repurposing in Glioblastoma: a computational and experimental workflow

**DOI:** 10.1101/2024.04.29.589520

**Authors:** Nazareno Gonzalez, Melanie Pérez Küper, Matías Garcia Fallit, Jorge A. Peña Agudelo, Alejandro J. Nicola Candia, Maicol Suarez Velandia, Guillermo A. Videla-Richardson, Marianela Candolfi

## Abstract

**Purpose:** Glioblastoma (GBM) remains a formidable challenge in oncology due to its invasiveness and resistance to treatment, i.e. surgery, radiotherapy, and chemotherapy with temozolomide. This study aimed to develop and validate an integrated model to predict the sensitivity of GBM to alternative chemotherapeutics and to identify novel candidate drugs and combinations for the treatment of GBM.

**Patients and Methods:** We utilized the drug sensitivity response data of 272 compounds from CancerRxTissue, a validated predictive model, to identify drugs with therapeutic potential for GBM. Using the IC50, we selected ’potentially effective’ drugs among those predicted to be blood-brain barrier permeable via *in silico* algorithms. We ultimately selected drugs with targets overexpressed and associated with worse prognosis in GBM for experimental *in vitro* validation.

**Results:** The workflow proposed predicted that GBM is more sensitive to Etoposide and Cisplatin, in comparison with Temozolomide, effects that were validated *in vitro* in a set of GBM cellular models. Using this workflow, we identified a set of 5 novel drugs to which GBM would exhibit high sensitivity and selected Daporinad, a blood-brain barrier permeant NAMPT inhibitor, for further preclinical *in vitro* evaluation, which aligned with the *in silico* prediction.

**Conclusion:** Our results suggest that this workflow could be useful to select potentially effective drugs and combinations for GBM, according to the molecular characteristics of the tumor. This comprehensive workflow, which integrates computational prowess with experimental validation, could constitute a simple tool for identifying and validating compounds with potential for drug reporpusing in GBM and other tumors.

## INTRODUCTION

Glioblastoma (GBM) is a tumor of glial origin that accounts for 50% of all primary malignant brain tumors in adults [1]. It is characterized by its invasiveness and intrinsic resistance to conventional therapy. In 2016, the WHO included distinctive genetic and epigenetic alterations to define various groups of gliomas [2–5]. The updated 2021 WHO classification of tumors of the central nervous system (CNS) [6] gives now a central role to the mutation status of the enzyme isocitrate dehydrogenase (IDH). Since then, three distinct molecular tumor categories have been designated for adult diffuse gliomas: Astrocytoma with mutated IDH (mIDH), Oligodendroglioma with mIDH and 1p/19q codeletion, and GBM with wildtype IDH (wtIDH). IDH-mutant diffuse astrocytic tumors are regarded as a unified entity, classified as CNS WHO grade 2, 3, or 4 based on various histological and molecular characteristics.

Standard therapy for GBM includes maximal safe surgical resection [7], adjuvant radiotherapy and chemotherapy with temozolomide (TMZ) [8, 9]. Patients with GBM face a poor median survival of 15 to 18 months, and approximately only 7% survive 5 years after diagnosis [1, 10]. Unfortunately, although the current treatment does not exert substantial clinical effects, it has remained unchanged since the introduction of TMZ in 2005 [11]. Consequently, there is an urgent need to explore novel therapeutic options to improve the treatment of GBM.

The Human Genome Project [12] has revolutionized cancer research by leveraging high-throughput data. Databases like the Gene Expression Omnibus (GEO) [13] and The Cancer Genome Atlas (TCGA) have facilitated the discovery of new biomarkers, enhancing cancer diagnosis, prognosis, and treatment. Variability in drug response among cancer patients is largely influenced by molecular tumor characteristics such as gene mutations, copy number variations, promoter methylation, and gene expression profiles. Bioinformatic methods, particularly differential gene expression studies, hold promise in predicting drug response [14, 15]. Utilizing gene expression data from cancer cell lines and drug sensitivity information, researchers aim to predict drug mechanisms and therapeutic potential in patients [16–18]. In line with this, Li et. al [19] developed an algorithm to predict IC50 values, in which aid to extrapolate drug sensitivity, and validated it using in vitro data from the Genomics of Drug Sensitivity in Cancer (GDSC) database. This model was then applied to predict IC50 values in over 10,000 tissue samples from TCGA. Using this algorithm, we developed a workflow that combines computational drug selection with in vitro validation to identify drugs with therapeutic potential for GBM., offering a transformative approach to identify therapeutic options and candidates for drug reporpusing in GBM to expedite the advance of translational neuro-oncology.

## RESULTS

### 1. Drug sensitivity prediction model

Our approach utilizes predicted IC50 values for 272 drugs from patient samples in TCGA obtained through CancerRxTissue [19] with *in silico* techniques, followed by evaluation in relevant GBM cellular models. Our workflow involves three stages (Fig. 1A): 1) identifying potential drugs; 2) *in silico* analyses; and 3) *in vitro* validation of predicted effects. In the first stage, data processing involves stratifying glioma patients based on WHO CNS classification and other characteristics to evaluate IC50 values, a critical parameter for understanding drug efficacy. We then select potentially effective drugs using the median of the medians of predicted IC50 values as a cut-off. The second stage consists of: 1) predicting blood-brain barrier (BBB) permeability; 2) evaluating target expression to ensure drug specificity; and 3) selecting relevant GBM cell lines based on target expression levels. The third stage involves experimental validation of predicted outcomes.

**Fig. 1:**
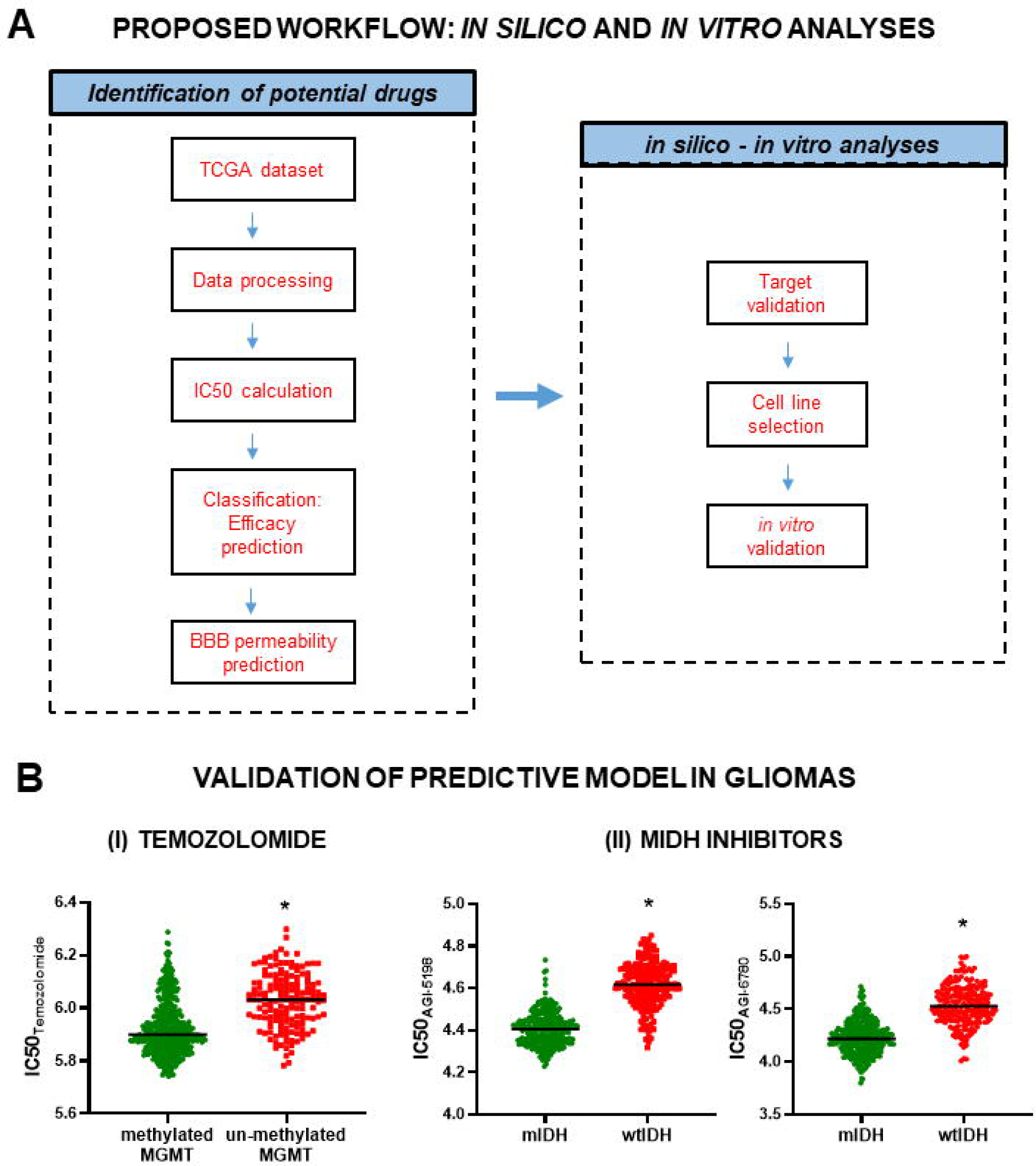
***Drug sensitivity prediction model*** **A)** The process involves data processing for the selected tumor type from TCGA datasets, IC50 estimation, efficacy prediction and subsequent classification. Then, *in-silico* evaluation for blood-brain barrier (BBB) permeability, target validation, and finally, the selection of cell lines to evaluate the selected drugs in relevant *in vitro* cancer models. **B)** To validate the algorithm, drugs that are already approved or tested for GBM treatment were evaluated using data from glioma biopsies deposited at TCGA: **I)** IC50 prediction for Temozolomide stratifying glioma patients according to the methylation of the promoter of the DNA repair enzyme MGMT (met, methylated; un-met, unmethylated); **II)** IC50 values for AGI-5198 and AGI-6780, specific inhibitors of mutated IDH enzyme (mIDH) in patients with mIDH and wild-type IDH (wtIDH). All IC50 values shown are ln (IC50) transformed.

To independently validate the prediction model developed by Li *et al.* [19], we first explored drugs commonly used in GBM. As expected, the algorithm predicts that treatment with TMZ will exert a stronger antitumoral effect (lower IC50) in glioma patients with methylation of the MGMT promoter, which leads to reduced expression of the DNA repairing enzyme (Fig. 1B I). In addition, the algorithm predicts that specific inhibitors of the mutated enzyme isocitrate dehydrogenase (mIDH) will exhibit a stronger effect (lower IC50) in patients with mIDH gliomas than in wtIDH glioma patients (Fig. 1B II). These findings suggest that the algorithm holds predictive potential for determining drug sensitivity in these patients.

### 2. Identification and experimental validation of chemotherapeutic drugs with therapeutic potential for GBM

After validating the potential utility of the predictive model, our goal was to identify alternative therapeutic strategies for adult gliomas among a pre-defined list of chemotherapeutic drugs (Fig. 2A). Initially, we assessed drug permeability across the blood-brain barrier (BBB) using various *in silico* tools [20, 21] (Fig. S1B). TMZ, carmustine, cyclophosphamide, fluorouracil, and cisplatin were predicted to cross the BBB, while contradictory results were obtained for Etoposide BBB permeability, our analyses predicted that GBM will exhibit high sensitivity to this drug. Moreover, Etoposide has been utilized for intracranial neoplasia in clinical settings [22]. While carmustine and TMZ exhibited the highest predicted IC50, i.e. lowest predicted antitumoral effect, Etoposide and Cisplatin emerged as potential drugs for which GBM would exhibit high sensitivity (Fig. 2A). Thus, we selected these drugs, as well as TMZ, the gold-standard for treatment in patients, for further *in silico* and *in vitro* validation. Comparing the predicted IC50 of these drugs between mIDH and wtIDH gliomas (Fig. 2B), we found that while wtIDH gliomas are predicted to exhibit a lower predicted response to TMZ and Cisplatin than mIDH gliomas, Etoposide would elicit a stronger effect (lower IC50) in wtIDH gliomas. Using neurospheres derived from mIDH and wIDH gliomas developed in mice by genetic engineering [23], we validated these predictions *in vitro* (Fig. 2B). Concentration-response curves in U251, LN229, and U87 cells showed that these cells were more sensitive to Etoposide followed by Cisplatin than to TMZ (Fig. 2C), aligning with the *in silico* prediction. To assess potential toxicity, we compared the cytotoxic effects of these drugs between murine normal astrocytes and GBM neurospheres (Fig. 2D). Normal astrocytes exhibited some sensitivity to TMZ, with GBM neurospheres showing very limited response. In contrast, Cisplatin and Etoposide demonstrated potent antitumor effects in this GBM model with no apparent toxicity in normal astrocytes (Fig. 2D). We selected Etoposide to further evaluate its therapeutic potential due to its superior antitumoral effect and favorable therapeutic window.

**Fig. 2:**
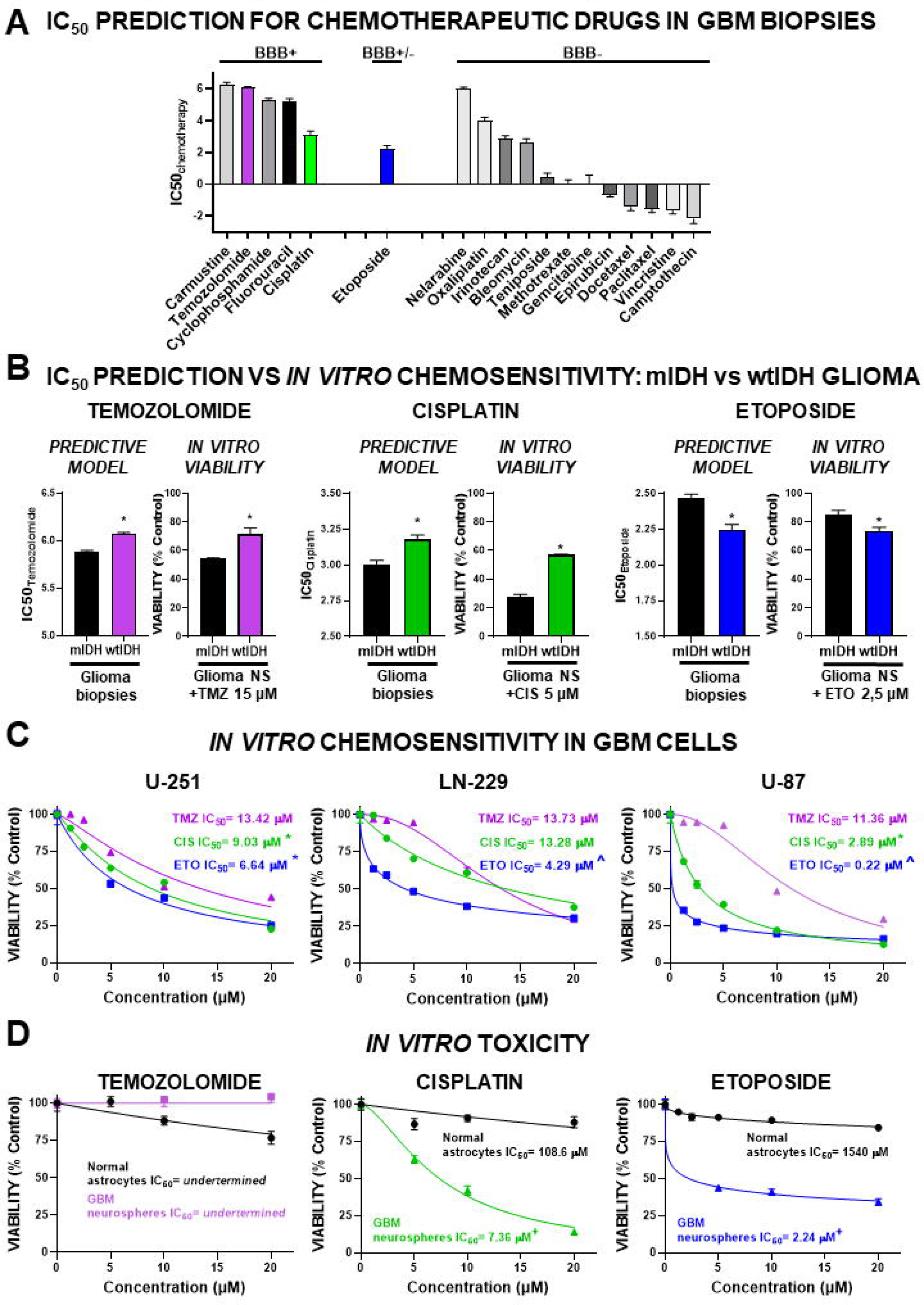
***In-silico IC50 prediction in glioma biopsies vs in vitro chemosensitivity in glioma cells: chemotherapeutic drugs*** **A)** IC50 prediction for chemotherapeutics drugs in mIDH and wtIDH glioma patients in comparison with IC50 for temozolomide, classified according to their blood-brain barrier (BBB) permeability prediction (BBB+, permeable; BBB+/-; ambiguous; BBB-; not-permeable). **B)** In-silico prediction and *in vitro* effect for temozolomide (TMZ), Cisplatin, Etoposide in mIDH and wtIDH glioma. Predicted IC50 values for these drugs in mIDH and wtIDH glioma patients (left panels). For *in vitro* validation, mIDH and wtIDH glioma neurospheres were treated at a fixed concentration of each drug (TMZ,15 µM; cisplatin, 5 µM; and Etoposide, 2.5 µM) for 72 hours and cell viability was determined by MTT assay. **C)** Commercial human GBM cell lines (U-251, LN229 and U-87 cells) and **D)** murine normal astrocytes and GBM neurospheres were treated with different concentrations of TMZ, Cisplatin, and Etoposide for 72 hours and cell viability was assessed by MTT assay. Concentration-response curves were plotted, and IC50 values were calculated through non-linear regression fit. *, p<0.05 vs. TMZ; ^, p<0.05 vs. all other drugs, ANOVA; +, p<0.05 vs. normal astrocytes; Student’s t test.

We found expression of DNA topoisomerase IIα (TOP2A), a target for Etoposide and other chemotherapy agents [24], in GBM sections from The Human Protein Atlas (THPA) [25] (Fig. 3A). Since our model predicted increased sensitivity to Etoposide in wtIDH than in mIDH gliomas, we compared the expression TOP2A across these glioma types and normal brain tissue. Our analysis revealed overexpression of TOP2A in wtIDH glioma biopsies compared to mIDH gliomas and normal brain (Fig. 3B). Treatment of mIDH (G01) and wtIDH (G08) glioma patient-derived cells with a fixed concentration of Etoposide confirmed higher sensitivity in wtIDH cells (Fig. 3D II).

**Fig. 3:**
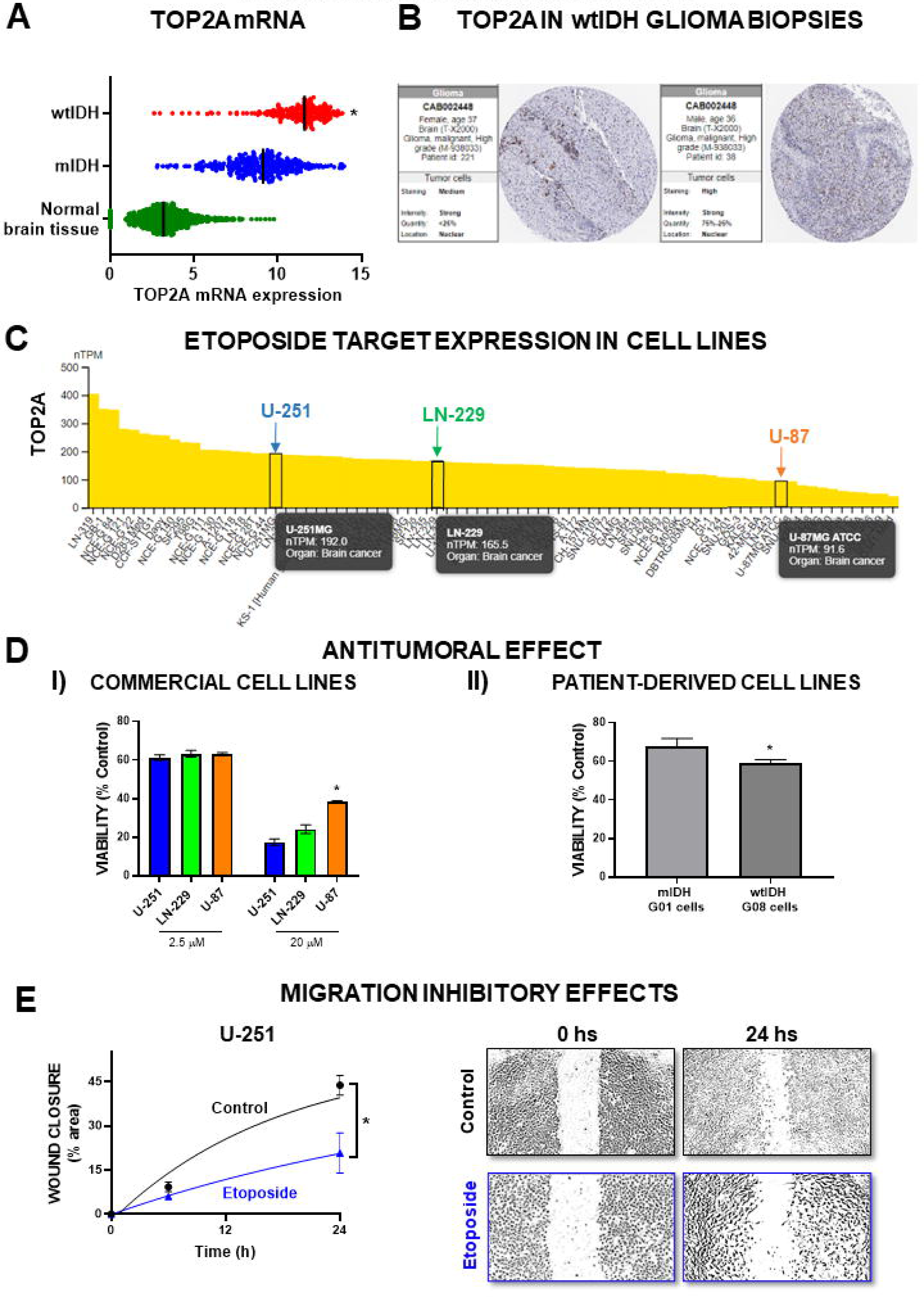
***In vitro preclinical evaluation of Etoposide in GBM cells*** **A)** TOP2A mRNA expression levels in normal brain tissue and in tumor biopsies from patients with mIDH or wtIDH gliomas. **B)** Representative IHC images with TOP2A expression in GBM biopsies extracted from The Human Protein Atlas. **C)** TOPA2 expression in human GBM cell lines using data from The Human Protein Atlas. **D) I)** U-251, LN-229 and U-87 GBM cells with different TOP2A expression levels were treated with different concentrations of Etoposide and viability was assessed by MTT assay after 72h. *, p<0.05 vs. U-251, ANOVA. **II)** Patient-derived mIDH and wtIDH glioma cells were treated with Etoposide (2.5 uM) to evaluate its cytotoxicity by MTT assay. *p<0.05, Student’s *t* test. **E)** U-251 GBM cells were seeded until reaching confluence and treated with Etoposide (1 µM) for 24 h. Cell migration was evaluated using the wound closure assay. *p<0.05 vs Control (nonlinear regression analysis).

Using information available at THPA [25] we found that the expression of TOP2A was higher in U251 cells, intermediate in LN229 cells and lower in U87 cells (Fig. 3C). We then assessed the sensitivity of these cell lines to Etoposide at high and low concentrations. As depicted in Fig. 3D I, 2.5 µM Etoposide demonstrated robust cytotoxicity across the three commercial cell lines, with the maximum response (20 µM) positively correlating with TOP2A expression levels. Analyzing mRNA expression levels of GBM biopsies, we observed a positive correlation between TOP2A and several epithelial-mesenchymal transition (EMT) markers (Fig. S3). To validate these findings, we used a wound closure assay, demonstrating that Etoposide inhibited U251 cell migration, a hallmark of EMT (Fig. 3E).

### 3. Identification of alternative drugs with therapeutic potential for GBM

Among all 272 drugs (Fig. 4AI, Table S1) we identified several BBB-permeable drugs (BBB+) (Fig. S2A) with high predicted sensitivity in GBM (low IC50) (Fig. 4AII, Table S2). We proceeded with the selection process of potential candidates for drug reporpusing based on rigorously defined criteria: target upregulation in the tumor compared to normal tissue (Fig. 4B); predicted higher efficacy compared to TMZ (Fig. 4C) and low reported toxicity in previous clinical trials for other diseases. Following these criteria, we compiled a shortlist of five drugs: Sepantronium bromide (specific BIRC5 inhibitor), Daporinad (specific NAMPT inhibitor), CUDC-101 (a potent inhibitor against HDAC, EGFR, and HER2 targets), HG6-64-1 (specific BRAF inhibitor), and QL-XII-47 (BMX and BTK inhibitor) (Fig. 4AII). Although all these drugs met the established parameters, only the target for Daporinad, nicotinamide phosphoribosyltransferase (NAMPT), was effectively present in GBM biopsies from IHC samples sourced from THPA (Fig. 4D). Moreover, overexpression of NAMPT was associated with worse prognosis in wtIDH glioma patients (Fig. 4E, Fig. S2B). Consequently, we selected Daporinad for preclinical *in vitro* assessment of therapeutic potential.

**Fig. 4:**
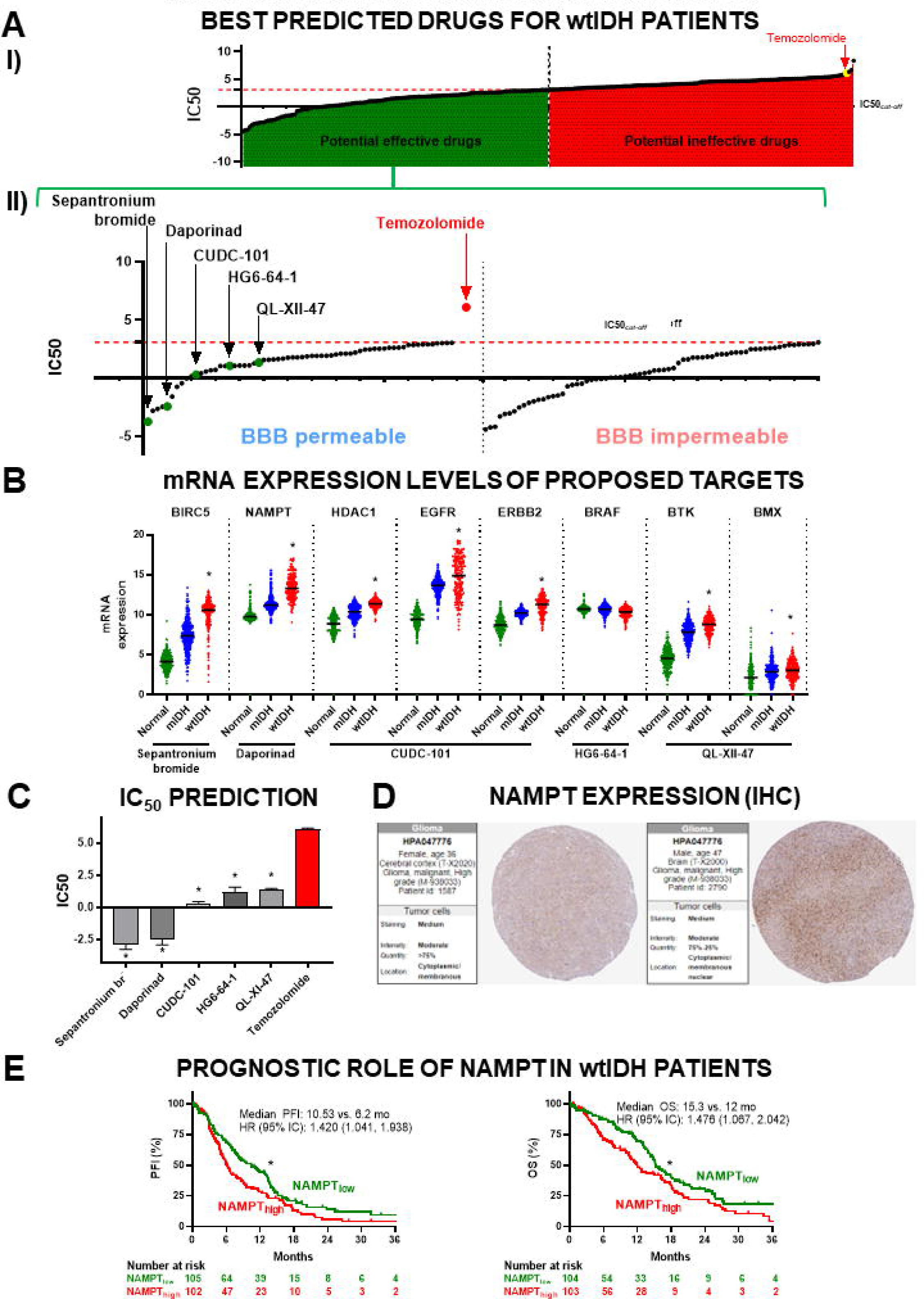
***Efficacy prediction model for alternative drugs with potential effect in GBM*** **A)** Drugs with the highest predicted efficacy in wtIDH patients, using **I)** the median of the median IC50 values for each of the 272 drugs to cut-off between potential “effective” and “ineffective”, and **II)** classifying the potentially effective drugs as BBB permeable (BBB+) or BBB not permeable (BBB-). According to pre-defined criteria a shortlist of 5 drugs was selected: Sepantronium bromide (specific BIRC5 inhibitor), Daporinad (specific NAMPT inhibitor), CUDC-101 (inhibitor against HDAC, EGFR and ERBB2), HG6-64-1 (specific BRAF inhibitor) and QL-XII-47 (BMX and BTK inhibitor). **B)** Evaluation of the mRNA expression levels of targets of the selected drugs in normal brain, and gliomas (miDH and wtIDH). **C)** IC50 prediction of the selected drugs in comparison with IC50 for TMZ for wtIDH patients. *, p<0.05, ANOVA. **D)** Representative IHC images with NAMPT expression in GBM biopsies extracted from The Human Protein Atlas. **E)** Progression-free-interval (PFI) and overall survival (OS) curves of wtIDH patients that were stratified according to NAMPT expression levels using the median of expression as a cut-off. Kaplan–Meier curves were created using data from UCSC Xena database and TCGA LGG-GBM cohorts. * p<0.05, Log-rank (Mantel-Cox) test. All IC50 values shown are ln (IC50) transformed.

Using the data from THPA we found expression of NAMPT in commercial GBM cell lines, which was as follows: U-87 cells: NAMPT^high^; U-251 cells: NAMPT^medium^; and LN-229 cells: NAMPT^low^ (Fig. 5A). Daporinad showed antitumoral effects in all these commercial cell lines, revealing a potent effect at remarkably low concentrations and showing that in conditions of higher NAMPT levels, such as in U87 cells, the effect of Daporinad was lower (Fig. 5B). This antitumoral effect was extended to cell cultures derived from GBM biopsies [26] (Fig. 5B). Additionally, Daporinad exhibited no evident effect on normal murine astrocytes while in murine GBM neurospheres its IC50 was comparable to that in human GBM cells (Fig. 5C).

**Fig. 5:**
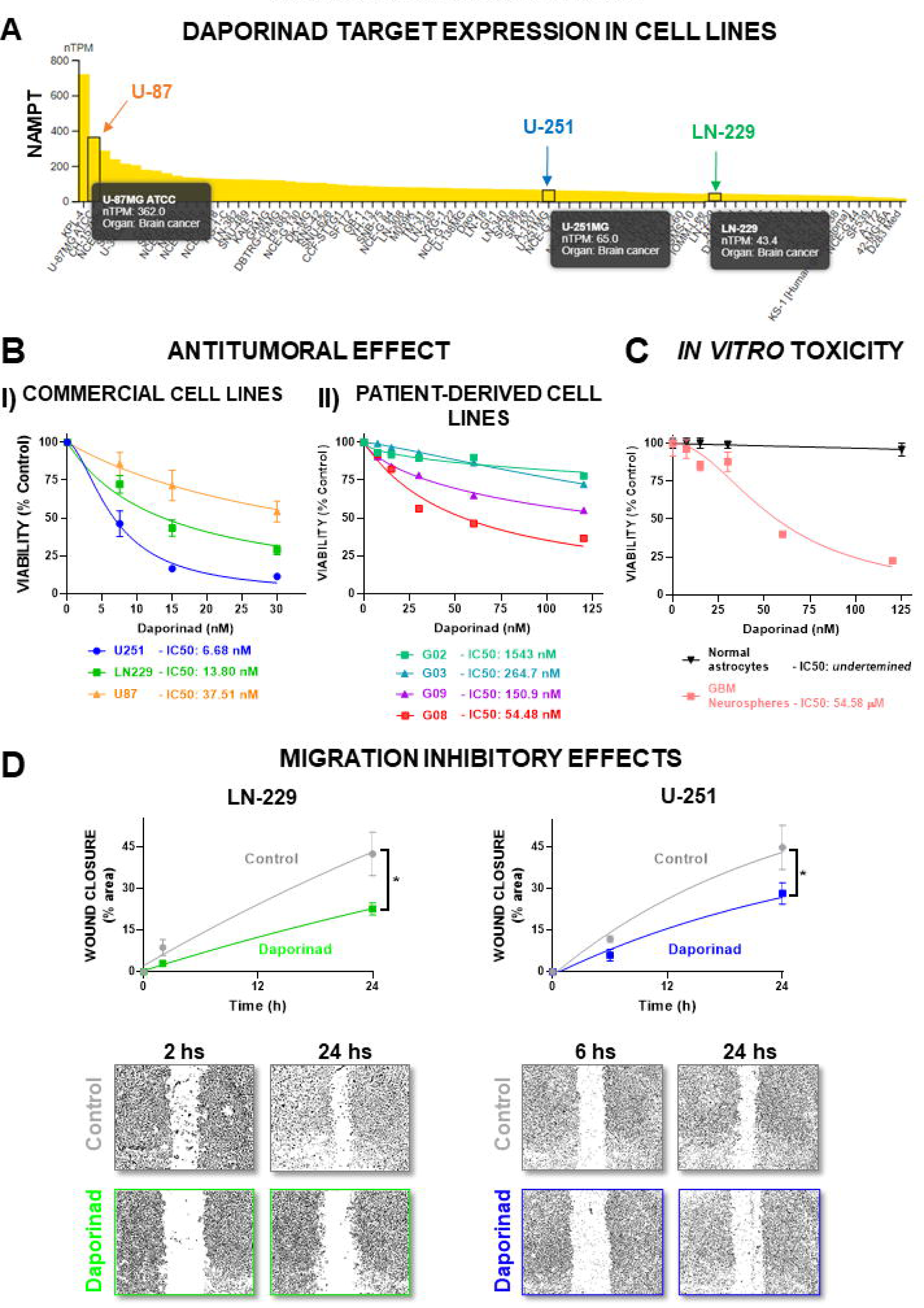
*In vitro* preclinical evaluation of Daporinad. **A)** Daporinad target (NAMPT) expression in human GBM cell lines, using data from The Human Protein Atlas. **B) I)** U-251, LN-229 and U-87 commercial GBM cells, **II)** cell cultures derived from GBM biopsies and **C)** murine normal astrocytes and GBM neurospheres, were treated with different concentrations of Daporinad for 72 h, and viability was assessed by the MTT assay. Dose- response curves were plotted, and IC50 values were calculated through non- linear regression fit**. D)** LN-229 and U-251 GBM cells were seeded until reaching confluence, treated with Daporinad (10 nM) for 24 h and migration was evaluated at different time points using the wound closure assay. *p<0.05 vs Control (nonlinear regression analysis).

Meta-analysis of GBM biopsies from TCGA revealed a positive correlation between NAMPT, and markers associated with EMT (Fig. S3). Thus, we evaluated the effect of Daporinad on the migration of GBM cells, finding that Daporinad effectively inhibited the migration of U251 and LN229 cells (Fig. 5D).

### 4. Identification of drug combinations with potential therapeutic efficacy in GBM

To identify potential drug combinations for GBM, we explored correlations between drug response and NAMPT mRNA expression levels in biopsies from GBM patients. We observed a positive correlation between TMZ ln(IC50) values and NAMPT expression levels (Fig. 6A), suggesting that patients with elevated tumor NAMPT levels will be less sensitive to TMZ. Lower predicted efficacy of a specific drug in patients with higher NAMPT levels suggests that its combination with Daporinad could enhance chemosensitivity. Consequently, we combined Daporinad with TMZ in patient derived GBM cell cultures with different NAMPT expression levels, *i.e.* NAMPT^high^ G02 GBM cells and NAMPT^low^ G09 GBM cells (Fig. 6B I). We treated these cells with Daporinad (60 nM), TMZ (150 μM) or both. This assay validated the predicted higher sensitivity of G09 NAMPT^low^ GBM cells to TMZ while, interestingly, TMZ increased the viability of NAMPT^high^ G02 GBM cells (Fig. 6B II). As shown above, Daporinad exerted a cytotoxic effect that was higher in NAMPT^low^ G09 cells than in NAMPT^high^ G02 GBM cells (Fig. 6B II). Tallying with the results from the predictive model, Daporinad sensitized both GBM cells to TMZ (Fig. 6B II).

**Fig. 6:**
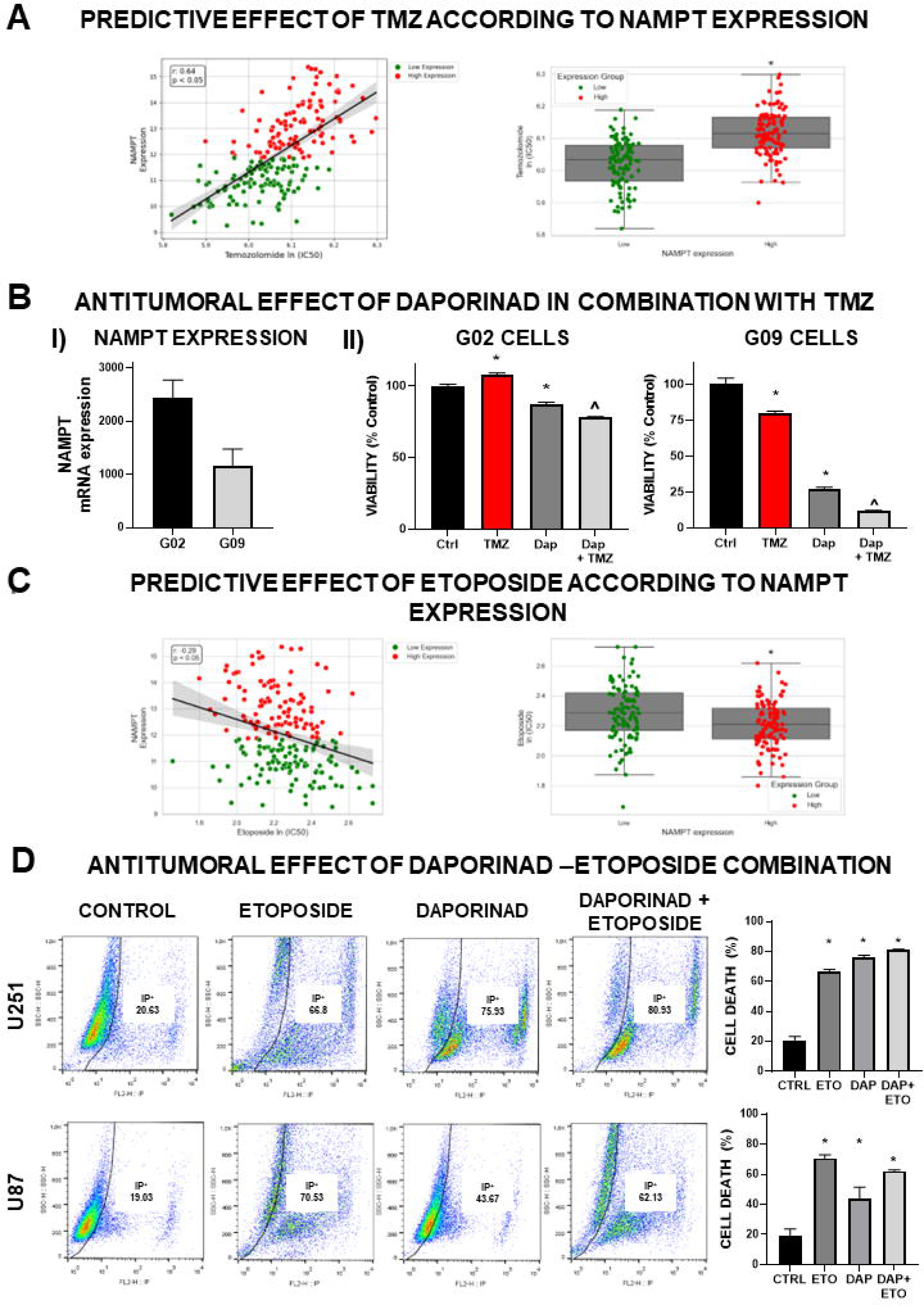
***Potential of Daporinad combination with chemotherapy in GBM cells*** **A)** Spearman correlation analysis between NAMPT expression in GBM patients and TMZ predictive effect (left panel; green: NAMPT low expression; red: NAMPT high expression). r, Spearman coefficient. Temozolomide predicted IC50 for wtIDH patients stratified according to NAMPT levels (right panel; green: NAMPT low expression; red: NAMPT high expression). wtIDH patients were classified according to NAMPT expression levels into “Low” and “High” expression, using the median of expression as a cut-off. * p <0.05; Mann- Whitney test. **B)** I) Relative expression levels of NAMPT analyzed by RNA-seq in G02 and G09 cells derived from GBM biopsies. G02 and G09 cells were incubated with Daporinad (60 nM) and Temozolomide (150uM) alone or in combination, and 72h later cell viability was assessed by MTT assay. **C)** Spearman correlation analysis between NAMPT expression in GBM patients and Etoposide predictive effect (left panel; green: NAMPT low expression; red: NAMPT high expression). r, Spearman coefficient. Etoposide predicted IC50 for wtIDH patients stratified according to NAMPT levels (right panel; green: NAMPT low expression; red: NAMPT high expression). wtIDH patients were classified according to NAMPT expression levels into “Low” and “High” expression, using the median of expression as a cut-off. * p <0.05; Mann-Whitney test. **D)** GBM cell lines (U-251 and U-87) were incubated with Daporinad (10 nM) and Etoposide (1 µM) alone or in combination and cell death was evaluated using the iodide propidium exclusion assay by FACS analysis. *p<0.05 vs control (One-way ANOVA).

Following the same rationale, we evaluated the predicted effect of Etoposide based on NAMPT expression. We identified a negative correlation between ln(IC50) values of Etoposide and NAMPT mRNA expression levels in GBM biopsies, suggesting that Etoposide would exert stronger antitumoral effects in patients with high NAMPT levels (Fig. 6C). Thus, it is possible that Daporinad might not be an ideal choice for combination with Etoposide. To validate this prediction, we assessed cell death by propidium iodide (PI) exclusion in NAMPT^high^ U-87 cells and NAMPT^low^ U-251 treated with Daporinad (10 nM), Etoposide (1 μM), or both. As we observed when assessing cell viability, Etoposide showed similar effects in both cell lines, while Daporinad exerted a stronger cytotoxic effect in U-251 cells in comparison with U-87 cells (Fig. 6D). In accordance with the prediction model, Daporinad did not improve the response of these cells to Etoposide (Fig. 6D).

Combining drugs with TMZ could be a promising approach to address GBM challenges, such as invasiveness and resistance to conventional therapies. We conducted an integrative analysis, assessing predicted sensitivity to TMZ and gene expression in canonical pathways [27] (Fig. S4, S5). TCGA GBM patients were classified into “low” and “high” expression groups for each pathway and predicted IC50 values for TMZ were evaluated in both groups. Pathways with higher TMZ IC50 values in the ’high’ expression group suggest that combining TMZ with drugs that inhibit those pathways could enhance its efficacy. This analysis revealed potential therapeutic combinations of TMZ with inhibitors of these pathways that were previously tested in clinical trials for other diseases (Table S2)

### 5. Validation of the predictive model in peripheral tumors

To explore whether this predictive model could also be applied to other tumors, we evaluated the predicted sensitivity of breast, prostate, and melanoma patients to drugs already used for the treatment of these tumors (Fig. 7). For breast cancer, the model predicts better effects for chemotherapeutics drugs as Docetaxel, Paclitaxel, Cisplatin and Epirubicin (a Doxorubicin derivative), in triple-negative breast cancer patients (basal-like subtype) in comparison with LumA-B and HER2 subtypes (Fig. 7A I). These results agree with clinical settings where chemotherapy is the standard treatment for triple—negative breast cancer tumors. Moreover, the model predicts that HER2-positive breast cancer patients will be more sensitive to specific HER2 inhibitors such as Lapatinib and Afatinib than Luminal (LumA-B) or Basal (triple negative) patients (Fig. 7A II). On the other hand, for prostate cancer the model predicts better effects for Bicalutamide, a specific androgen receptor (AR) inhibitor indicated for metastatic prostate cancer, in patients with higher AR expression levels and increasing Gleason scores, which indicates a higher probability of dissemination (Fig. 7B III). Lastly, the model predicts that Dabrafenib, a specific inhibitor for the treatment of cancers associated with a mutated version of BRAF, will show higher effect (lower IC50) in melanoma patients with this mutation (Fig. 7C IV). We observed similar results for Trametinib, another mutated BRAF inhibitor (Fig. 7C V). In summary, our results show that the predictive model could effectively predict the sensitivity of different types of tumors to a large set of novel and traditional drugs.

**Fig. 7:**
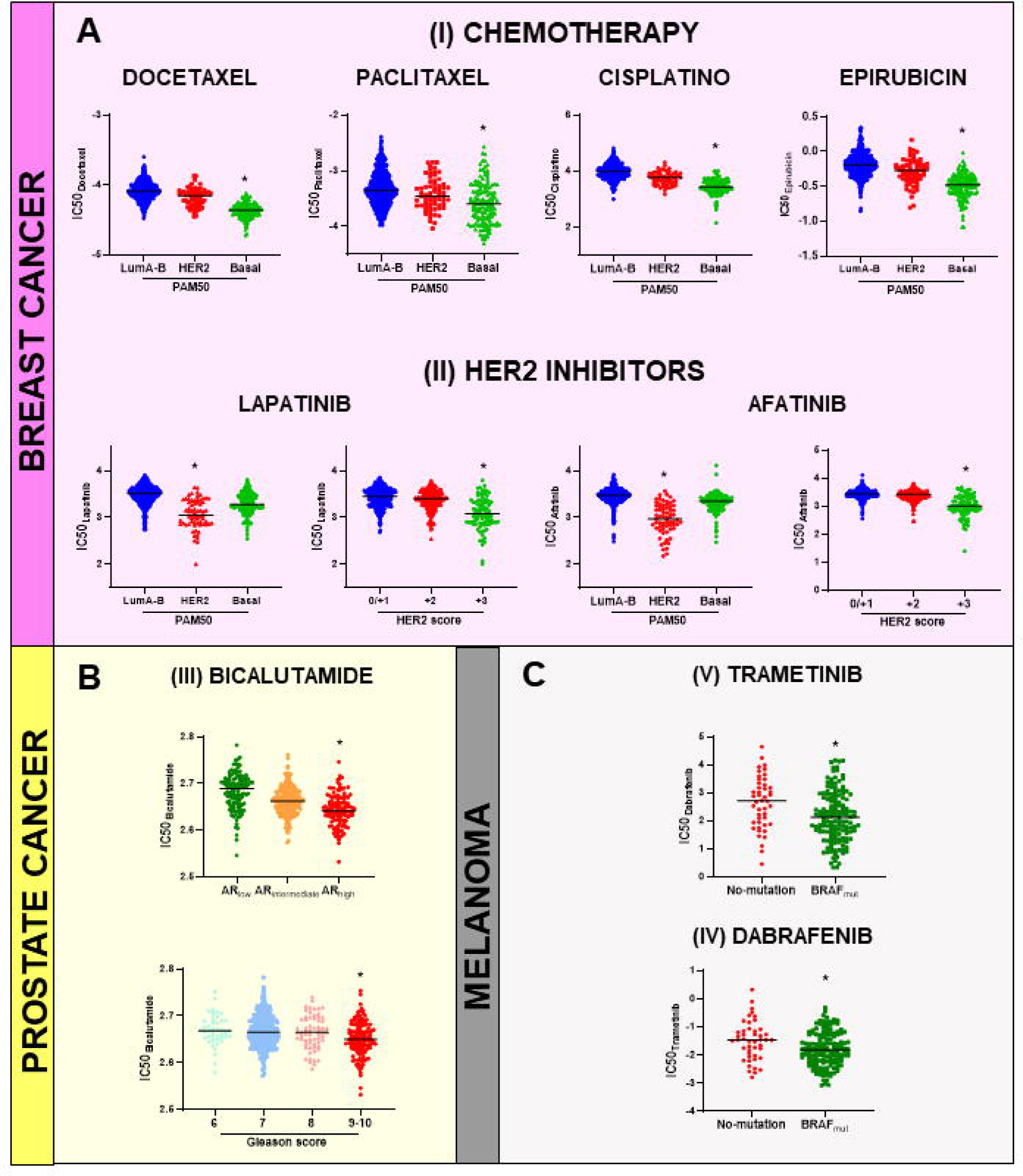
***Validation of the predictive model in peripheral tumors*** We evaluated different predictive IC50 values for drugs that are already approved or tested for other types of cancer, such as breast, prostate, and melanoma. **A) I)** Docetaxel, Paclitaxel, Cisplatin and Epirubicin (a Doxorubicin derivative) will show better effects in triple-negative breast cancer patients (basal-like subtype). **II)** Specific HER2 inhibitors such as Lapatinib and Afatinib will show better effects in HER2-positive breast cancer patients. **B) III)** Bicalutamide, a specific androgen receptor (AR) inhibitor indicated for metastatic prostate cancer, will show better effect in prostate cancer patients with higher AR expression levels and increasing Gleason scores (higher Gleason score indicates a higher probability of dissemination). **C) IV)** Dabrafenib, a specific inhibitor for the treatment of cancers associated with a mutated version of BRAF, will show higher effect (lower IC50) in melanoma patients with BRAF mutation. **V)** As observed in clinical settings, Trametinib will show better effect (lower IC50) in melanoma patients with BRAF mutation. *, p<0.05; Mann-Whitney test or ANOVA. All IC50 values shown are ln (IC50) transformed.

## DISCUSSION

Although the median survival of GBM patients is only 15-18 months [10], the standard treatment for these patients has remained unchanged for almost 20 years [11]. For this reason, it is necessary to explore alternative options to enhance the therapeutic response of GBM cells. In this study we developed an integrated workflow to predict the sensitivity of GBM to novel candidate compounds to expand the drug portfolio to treat GBM. The overall aim of our research was to offer a simple tool for identifying and validating potential drugs and therapeutic combinations to be further explored to treat adult GBM. The selection is based on the prediction of drug efficacy and permeability across the blood-brain barrier. This workflow allows to integrate a myriad of available resources (gene expression databases, clinical information of patients, *in-silico* tools, and software applications) to facilitate drug development or selection.

In the era of personalized therapy, GBM seems to be fitted to this kind of approach. However, until now, the use of personalized genomic medicine has been unsuccessful, in part, because the efforts were put into the development of novel treatment options that target unique features of each patient as single agents. It is very unlikely to define a unique identity of GBM, given its high genomic heterogeneity (inter- and intra-tumor, spatial and temporal) [28, 29]. Novel treatments have shown promising preclinical effects [30–34], but the dramatic scenario of patients with GBM needs the development of faster, safer, and novel approaches. Here is where the reporpusing of well-known drugs represents an alternative for these patients. They are safer, more cost-effective, and can move from the laboratory to clinical use more quickly.

Taking the lead of Li et. al [19], our strategy involves using the predicted effects of more than 270 drugs to propose novel therapeutic strategies for GBM. We validated the *in-silico* predictions by assessing the effects of the selected drugs in relevant cellular models. Initially, we used the prediction model to evaluate the therapeutic potential of different well-known chemotherapeutic drugs as alternative options to TMZ. Among the BBB+ cohort, we found that many drugs that have been used as salvage therapy for TMZ-refractory recurrent gliomas, such as cyclophosphamide, fluorouracil, and cisplatin, exhibit higher predicted efficacy than TMZ in GBM patients. Despite the model was not conclusive about the ability of Etoposide to cross the BBB, this drug has been used in GBM patients alone or in combination with TMZ, cisplatin and other drugs [22, 35–37]. We observed that Etoposide not only showed higher predicted and *in vitro* anti- cancer effects than TMZ, but also it showed very low toxicity in normal cells. Moreover, the lower expression levels of TOP2A in normal brain tissue in comparison with glioma tumors anticipated the lack of toxicity of this drug in non-neoplastic cells. In addition, amid the three drugs tested, our model predicted higher sensitivity in GBM than mIDH gliomas only for Etoposide, which was validated experimentally using murine glioma neurospheres with a specific genetic background [23]. Approved for anticancer therapies for almost 40 years [38, 39], Etoposide is used in the management and treatment of various cancers, such as refractory testicular tumors, and as a first-line treatment in combination with cisplatin for small-cell lung cancer [40]. Consistent with our findings, this drug has demonstrated comparable effects in gliomas [41]. The proposed use of Etoposide for GBM is not novel. Several preclinical studies have demonstrated that TOP2A inhibitors could be effective in the treatment of GBM [42, 43]. Moreover, several clinical trials have been conducted to evaluate its effectiveness and utilization for GBM [44–53]. Thus, improvement of the local availability of this drug in the tumor could improve its therapeutic efficacy.

Irinotecan, methotrexate, vincristine, or paclitaxel were predicted here to be more effective than TMZ, Cisplatin or Etoposide. However, they are expected not to cross the BBB. In fact, they are extensively used in chemotherapy and several CNS pathologies but have shown limited efficacy in GBM [54]. Although overcoming the BBB and achieving effective targeting with therapeutic compounds pose significant challenges, these drugs could be subjected to drug delivery approaches to overcome the BBB in gliomas. Although they have shown limited efficacy in GBM patients [54], numerous invasive and non-invasive methods that aim to improve drug biodistribution in gliomas [55] could enhance their efficacy.

We defined several criteria for selecting novel potential candidates for GBM, based on favorable toxicity profiles, druggability and prognostic role of its targets. We identified numerous drugs with therapeutic potential for GBM and among non-traditional compounds we established a shortlist of 5 drugs with different mechanisms of action. This first selection includes to Sepantronium bromide (specific BIRC5 inhibitor); Daporinad (specific NAMPT inhibitor); CUDC-101 (a potent inhibitor against HDAC, EGFR, and HER2 targets); HG6- 64-1 (specific BRAF inhibitor); and QL-XII-47 (BMX and BTK inhibitor). We focused on Daporinad, also known as APO866 or FK866, which has shown antitumor effects in different types of cancer [56–60] and has been tested in several clinical trials for Chronic Lymphocytic Leukemia B cells [61], Melanoma [62], and Cutaneous T-cell Lymphoma [63], with good safety profiles. Our results showed that its target, NAMPT, was not only overexpressed in gliomas in comparison with normal brain tissue, but also showed higher expression levels in wtIDH than in mIDH glioma biopsies. Furthermore, we found that high levels of NAMPT were significantly correlated with a worse prognosis in wtIDH patients. Experimental evaluation validated these predictions showing anti-tumoral effects of Daporinad in several GBM cellular models without apparent toxicity in normal cells.

NAMPT is a rate-limiting enzyme involved in the biosynthesis of nicotinamide adenine dinucleotide (NAD+), a molecule that regulates multiple metabolic processes in normal and neoplastic cells [64, 65], which exhibits a high demand for NAD+ [66]. Exacerbation of the NAD+ pathway is a hallmark in GBM [67] and NAMPT has been proposed to play a central role in the maintenance of glioma stem cell phenotype [68]. NAMPT has been also involved in the radio-resistance of GBM cells. Transfer of NAMPT via microvesicles (MVs) has been proposed to disseminate radio-resistance within the glioma tumor microenvironment [69].

Although Daporinad displayed promising preclinical outcomes, it lacked consistent tumor response in clinical trials for chronic lymphocytic leukemia B cells [61], melanoma [62], and cutaneous T-cell lymphoma [63]. Efforts to widen the therapeutic window and enhance NAMPT inhibitors’ efficacy include identifying predictive response biomarkers and developing second-generation NAMPT inhibitors [70]. Furthermore, recent research highlighted the potential interplay between the microbiota and the efficacy of NAMPT inhibitors [71, 72]. Gut microorganisms can mitigate the cytotoxic effects of Daporinad by boosting NAD production through nicotinamidase activity, facilitating nicotinamide conversion into nicotinic acid, a precursor in the alternative deamidated NAD salvage pathway [71]. Thus down-modulating gut microbiota with antibiotics has been proposed to optimize the antitumoral effects of this NAMPT inhibitor [72].

We also investigated the predictive capabilities of our workflow in defining candidate drug combination strategies, offering fresh insights into enhancing therapeutic efficacy. Our findings suggest that wtIDH tumors with low NAMPT expression may exhibit greater sensitivity to TMZ, and experimental validation revealed that NAMPT inhibition with Daporinad enhanced the response to TMZ in patient-derived cells refractory to TMZ. These results are consistent with findings demonstrating that NAMPT inhibitors sensitize human GBM commercial cell lines to TMZ treatment [73]. Likewise, the correlations derived from this analysis could serve as indicators of TMZ response and the molecular characteristics of specific pathways. We found potential combinations of TMZ with several inhibitors of canonical pathways, such as the ALK pathway, CTLA4 pathway, HDAC targets, HER2 amplified, PI3K cascade, RB pathway, and retinol metabolism. In fact, several of these potential effective combinations of TMZ with specific inhibitors are currently being tested in pre-clinical and clinical trials. For instance, Crizotinib, an ALK inhibitor, has shown promise in combination with Temozolomide [74–76]. Panobinostat, a non-selective histone deacetylase inhibitor, enhances the effect of TMZ [77, 78], while Lapatinib, a dual tyrosine kinase inhibitor of HER2 and EGFR pathways, has been tested in clinical trials for newly-diagnosed GBM [79] This approach aligns with the concept of combination therapy, aiming to target multiple molecular pathways simultaneously for a more comprehensive and effective treatment strategy [80–82] . Ultimately, this integrative analysis serves as a hypothesis-generating tool, proposing novel therapeutic combinations with the potential for improved efficacy in cancer treatment.

We recognize the necessity of assessing the combination of chosen drugs with TMZ. Despite its limited efficacy, TMZ has remained the gold standard for GBM treatment for almost two decades. While it represents a critical advancement in the field when introduced in 2005, it is an unfortunate reality that its clinical impact has not met the expectations initially set. Combining selected drugs with TMZ provides a compelling approach to tackle GBM challenges, such as its invasiveness and intrinsic resistance to conventional therapies. Therefore, demonstrating their efficacy in combination with TMZ offers a pathway for accelerated approval, facilitating the efficient translation of our findings into clinical practice.

Considering that TCGA comprises 10 thousand samples from 32 tumor types providing access to mutation status, copy number alterations, promoter methylation, and gene expression profiles, our workflow could constitute a simple and versatile tool for selecting optimal treatments from a group of 272 drugs, most of which are already approved for clinical use in other indications, offering widespread applicability across diverse patient populations based on the molecular diagnosis of their tumors.

## CONCLUSION

In conclusion, our proposed workflow may enable the identification of pharmacological strategies with therapeutic potential for GBM, a pathology without improvements in its treatment for the past two decades. The selection of drugs that have already proved to be non-toxic in patients with other diseases could expedite the translation of these strategies into clinical practice.

## MATERIALS AND METHODS

### Drug prediction workflow

We utilized clinical and transcriptomic data from The Cancer Genome Atlas (TCGA) dataset as a primary resource for identifying potential drugs for GBM. Predictive IC50 values of 272 drugs on TCGA glioma patients were obtained from CancerRxTissue [19], a parameter used as a quantitative measure for drug efficacy. To predict the potential efficacy of identified drugs we utilized the median of the medians of the IC50 values to facilitate the selection of candidates for further investigation. In-silico analyses were conducted to predict the permeability of identified drugs through the Blood-Brain Barrier (BBB) and to validate the targets of identified drugs, based on pre-defined criteria, such as predicted responsiveness for GBM; upregulation of expression levels of the target in the tumor compared to normal tissue; the ability to cross the blood-brain barrier; and a predicted higher efficacy compared to TMZ. Then, strategic cell line selection was done by analyzing mRNA expression levels of the targets using data from The Human Protein Atlas [83]. Selected drugs were subjected to *in vitro* validation (describe below).

### Glioma biopsies datasets

Clinical, genomic, and transcriptomic data of glioma patients from the TCGA LGG-GBM cohort and transcriptomic data from normal samples were obtained from GTEx dataset. We stratified patients according to the mutational status of IDH1/2 into two groups: mIDH and wtIDH patients. Expression data of TOP2A, NAMPT and canonical targetable signaling pathways data were downloaded from https://tcga-data.nci.nih.gov/ via Xena Browser developed by UCSC [84].

### Peripheral tumors biopsies datasets

Clinical, genomic, and transcriptomic data of breast cancer, prostate cancer and melanoma patients were obtained from the TCGA BRCA, TCGA PRAD and TCGA SKCM, respectively.

### Potential combination analyses of Temozolomide/Daporinad with other drugs

We conducted an integrative analysis of Temozolomide/Daporinad response and gene expression pathways using Python with the Pandas, Matplotlib, and Seaborn libraries. For each drug-pathway combination, we generated scatter plots with linear regression lines. Furthermore, we created boxplots with stripplots to illustrate the distribution of drug response in low and high pathway expression groups. Samples were classified in two groups, using median expression values as *cut-off*.

### Drugs

Daporinad (also known as FK866 or APO866), a competitive inhibitor of nicotinamide phosphoribosyltransferase (NMPRTase; NAMPT), was obtained from Selleck Chemicals (Cat# S2799). The compound was dissolved in dimethyl sulfoxide (DMSO) to achieve a final concentration of 10 mM. Aliquots of the stock solution were prepared to prevent repeated freeze–thaw cycles and stored at -80 °C until later use in experiments. Cisplatin and Etoposide was obtained from Microsules Argentina (Buenos Aires, Argentina). Temozolomide (TMZ) was obtained from Sigma (St. Louis,MO, USA).

Dulbecco’s Modified Eagle Medium (DMEM; Cat# 12100046), Dulbecco’s Modified Eagle Medium: F-12 Nutrient Mix (DMEM/F-12; Cat# 12500062), Neurobasal Medium (Cat# 21103049), B-27 and N-2 supplements (Cat# A35828-01 and Cat# 17502-048, respectively), Geltrex LDEV-Free Reduced Growth Factor Basement Membrane Matrix (Cat# A14132-02), penicillin– streptomycin (Cat# 15140122), trypsin–EDTA (0.025%, Cat# 25200114) were obtained from Gibco (Invitrogen, Carlsbad, CA, USA). Fetal bovine serum (FBS) was acquired from Natocor (Cordoba, Argentina).

### Cell Culture

Human GBM commercial cell lines (U-251, LN-229 and U-87), and GBM neurospheres derived from murine (wtIDH and mIDH) gliomas were kindly donated by Dr Maria G Castro (University of Michigan School of Medicine, Ann Arbor, MI, USA). Human GBM commercial cell lines were maintained routinely in Dulbecco’s Modified Eagle Medium (DMEM) (Sigma, New York, NY, USA), supplemented with 5% fetal bovine serum (FBS) and 1% penicillin-streptomycin (PS) (Sigma, New York, NY, USA), pHL7.4, under 5% CO2 atmosphere and 37°C. Once the cells reached 80% of confluence, they were dissociated with 0.05% trypsin-ethylenediamine tetraacetic acid (EDTA) and subcultured in 100mm plastic petri dishes every three days. Murine GBM neurospheres were cultured in DMEM-F12 supplemented with 1% penicillin-streptomycin, 1X B-27, 1X N-2, 100 g/mL Normocin, 20 ng/mL bFGF, and 20 ng/mL EGF. Neurospheres were collected and disaggregated using accutase. Patient-derived gliomas stem cells used in this study were previously isolated from human biopsies following relevant guidelines and national regulations. Cell lines, named G02, G03, G08 and G09, have been described previously [26]. The use of these cultures for biomedical research was approved by the Research Ethics Committee “Comité de Ética en Investigaciones Biomédicas de la Fundación para la Lucha contra Enfermedades Neurológicas de la Infancia (FLENI)”. These cells were cultured on Geltrex-coated Petri dishes with serum-free neurobasal medium supplemented with glucose, sodium pyruvate, PBS-BSA (7.5 mg/mL), 1X B27, 1X N2, 20 ng/mL bFGF and EGF, 2 mM L-glutamine, 2 mM non-essential amino acids, and 50 U/mL penicillin/streptomycin. Cells were harvested using accutase.

### Cell Viability

The anti-tumoral effects of chemotherapeutic drugs (Temozolomide, Cisplatin and Etoposide) and Daporinad were assessed using a 3-(4,5-dimethylthiazol-2- yl)-2,5-diphenyltetrazolium bromide (MTT) colorimetric assay. For this analysis, 5,000 cells were seeded per well in 96-well plates. After 24 hours, cells were washed and incubated with 100 μL of the respective treatments (diluted in the previously described culture medium). Following 72 hours of incubation, the treatment was removed, and the wells were washed. Subsequently, 110 μL of MTT 450 μg/mL (Molecular Probes, Invitrogen, Thermo Fisher Scientific,Waltham,MA, USA) in Krebs–Henseleit solution was added to each well. Plates were then incubated for 4 hours at 37°C, and after this incubation period, were analyzed on a spectrophotometer by measuring the absorbance at 595 nm. Dose-response curves were plotted, and IC50 values were calculated through non-linear regression fit using GraphPad Prism software.

### Wound closure assay

To assess the migration of human glioblastoma multiforme (GBM), U-251 and LN-229 cells in response to Etoposide and Daporinad, 100.000 cells were seeded per well in 24-well plates under culture conditions described before. After 24 hours, cells were incubated with or without Etoposide (1 µM) or Daporinad (10 nM) for 24 hours. Subsequently, a wound was made by carefully scratching the confluent cell culture using a micropipette tip. After the scratch, cells were thoroughly washed with phosphate-buffered saline (PBS) and then reincubated with or without Etoposide/Daporinad in complete Dulbecco’s Modified Eagle Medium (DMEM) without serum. Finally, cells were photographed at various time points, and wound area was quantified using ImageJ Software.

### Propidium Iodide Exclusion Assay

Quantification of cell death (apoptosis) was conducted using propidium iodide staining followed by FACS analysis. A total of 60,000 cells were seeded per well in 24-well plates and after 24 hours, they were treated with DMSO, Etoposide (1 µM), Daporinad (10 nM) or combination of both. Following 72 hours of treatment, cells were washed with PBS and dissociated using 0.025% trypsin– EDTA. Supernatants and detached cells were collected from each treatment. Samples were centrifuged for 5 min at 1500 rpm and the supernatant was discarded. For the preparation of the propidium iodide (PI) stock solution, 1 mg of PI was dissolved in 1 mL of distilled water; then, the working solution was prepared using 1 μL of the stock solution in 100 μL of PBS. Cells were resuspended with 200 μL of the working solution and immediately analyzed using a BD FACSCalibur Flow Cytometer (BD Biosciences). Twenty thousand events were then analyzed by flow cytometer on the fluorochrome PI channel (488 nm excitation laser and long pass filter 556 and band pass filter 616/23 for emission). Data analysis was performed using FlowJo™ v10 Software (BD Biosciences).

### Statistical analyses

Statistical analyses were performed using GraphPad Prism version 8 software (GraphPad Software). Data normality was assessed using the Kolmogorov-Smirnov test prior to conducting parametric statistical tests. Continuous variables were compared using Student’s *t*-test or one-way analysis of variance (ANOVA). Correlations between continuous variables were evaluated using Spearman correlation analysis. Kaplan-Meier curves were analyzed using Log-rank test. Differences were considered significant when p-value < 0.05.

## ACKNOWLEDGEMENTS

This work was supported by Consejo Nacional de Investigaciones Científicas y Técnicas (CONICET, Fellowships to A.J.N.C, J.A.P.A. and M.G.F.); Instituto Nacional del Cáncer (Asistencia Financiera a Proyectos de Investigación en Cáncer IV to M.C.); Agencia Nacional de Promoción Científica y Tecnológica (PICT-2018-3088, PICT- 2019-00117 and PICT-2022-11-00074 to M.C; fellowship to A.J.N.C, M.P.K. and N.G); Fundación Florencio Fiorini (2023 Research grant to N.G. and 2023 Cancer Research Award to M.C) and Consejo Interuniversitario Nacional (fellowship to M.G.F.).

## CONTRIBUTIONS

**Conceptualization**: M.C; N.G; M.P.K. **Methodology**: M.C; N.G; M.P.K. **Validation**, M.C; N.G; M.P.K. **Formal analysis**, M.C; N.G; M.P.K.; V.R.G.A **Investigation**, N.G; M.P.K; M.G.F; J.A.P.A.; A.J.N.C **Resources**, M.C. **Writing—original draft preparation**, N.G; M.P.K and M.C. **Writing—review and editing**, N.G; M.P.K. and M.C. **Visualization**; N.G and M.C. **Supervision**, M.C. **Project administration**, M.C. **Funding acquisition**, N.G and M.C. All authors have read and agreed to the published version of the manuscript.

## SUPPLEMENTARY FIGURES

**Fig. S1: *TOP2A correlations with EMT gene expression*** Spearman correlation between TOP2A expression levels and proteins involved in epithelial-mesenchymal transition (EMT). r, Spearman coefficient. *, p<0.05.

**Fig. S2: *Criteria for the selection of potentially effective drugs for GBM.* A)** Blood-brain barrier (BBB) permeability prediction of drugs evaluated in this study. BBB+, permeable; BBB-; not-permeable. **B)** Forest plot of progression-free interval (PFI) and overall survival (OS) in wtIDH patients between low and high expression levels of the different targets for the selected drugs.

**Fig. S3: *NAMPT correlations with EMT gene expression.*** Spearman correlation between NAMPT expression levels and proteins involved in epithelial-mesenchymal transition (EMT). r, Spearman coefficient. *, p<0.05.

**Fig. S4: *Predictive effect of Temozolomide according to the expression of targets for alternative drugs.*** Temozolomide predicted IC50 for wtIDH patients classified according to canonical pathways expression levels into “Low” (green) and “High” (red) expression, using the median of expression of each pathway as a cut-off. * p <0.05; Mann-Whitney test.

**Fig. S5: *Correlations of Temozolomide predictive effect with the expression of targets for alternative drugs.*** Spearman correlation between Temozolomide predicted IC50 and expression levels of specific canonical pathways. Each point represents a sample, color-coded by the expression level of the pathway (green: Low, red: High). Black lines indicate linear regression. r, Spearman coefficient.

**Fig. S6: *Predictive effect of Daporinad according to the expression of targets for alternative drugs.*** Daporinad predicted IC50 for wtIDH patients classified according to canonical pathways expression levels into “Low” (green) and “High” (red) expression, using the median of expression of each pathway as a cut-off. * p <0.05; Mann-Whitney test.

**Fig. S7: *Correlations of Daporinad predictive effect with the expression of targets for alternative drugs.*** Spearman correlation between Temozolomide predicted IC50 and expression levels of specific canonical pathways. Each point represents a sample, color-coded by the expression level of the pathway (green: Low, red: High). Black lines indicate linear regression. r, Spearman coefficient.

**Table S1: Ranking of drug efficacy prediction for novel drugs with potential effects in GBM.**

**Table S2: Ranking of drug efficacy prediction model for BBB^+^ and BBB^-^ drugs with potential effects in GBM.**

**Table S3: Summary of Temozolomide correlations with canonical pathways and potential drug combinations for GBM.**

**Table S4: Summary of Daporinad correlations with canonical pathways and potential drug combinations for GBM.**

## Notes

### Competing Interest Statement

The authors have declared no competing interest.

### Summary of Updates

We need to perform a modification of the title.

