## Supplementary Figures for "Identification and validation of drugs for repurposing in Glioblastoma: a computational and experimental workflow"

*Fig. S1: TOP2A correlations with EMT gene expression*

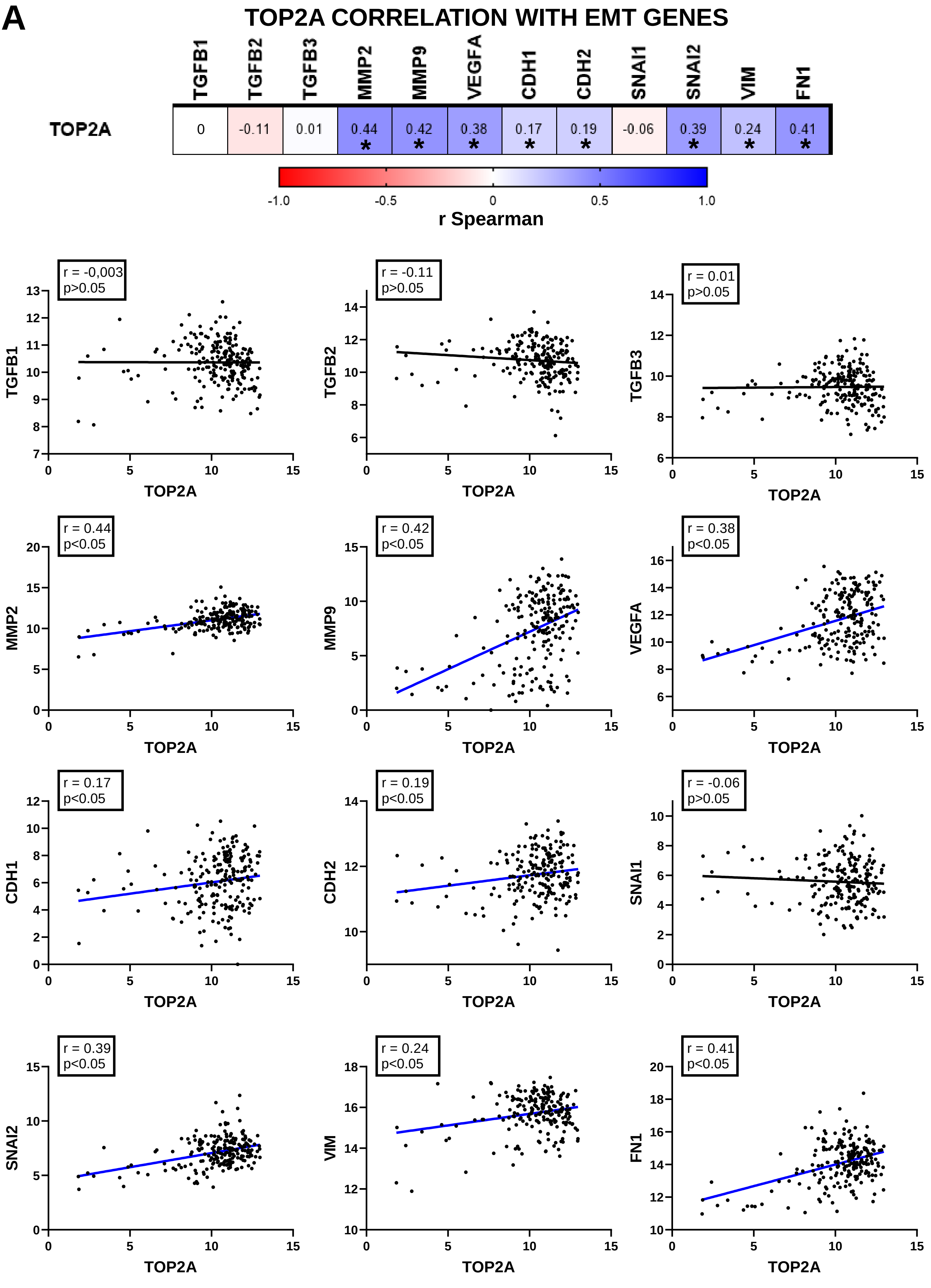

*Fig. S2: Criteria of selection of known drugs for GBM*

## A

### *BBB* PERMEABILITY *PREDICTION*

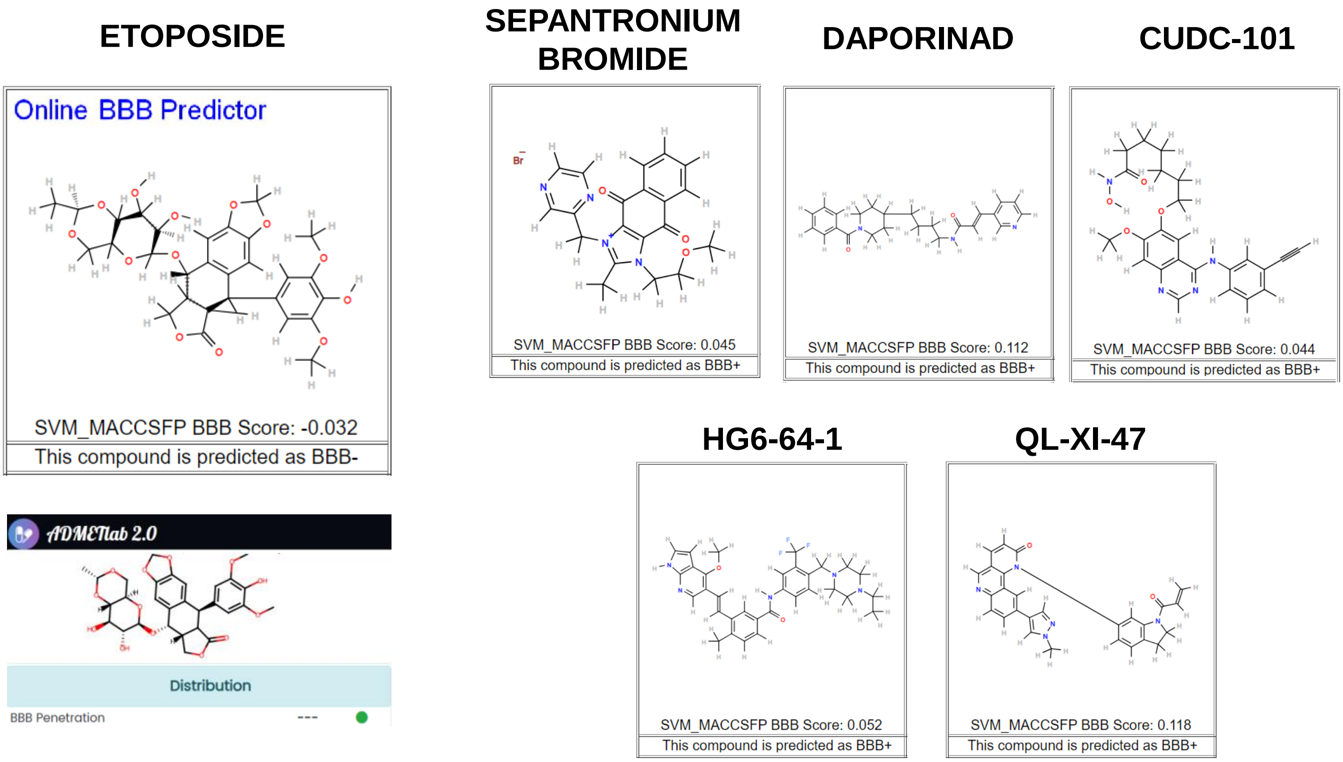


## B

### PROGNOSTIC ROLES OF THE TARGETS OF THE SELECTED DRUGS

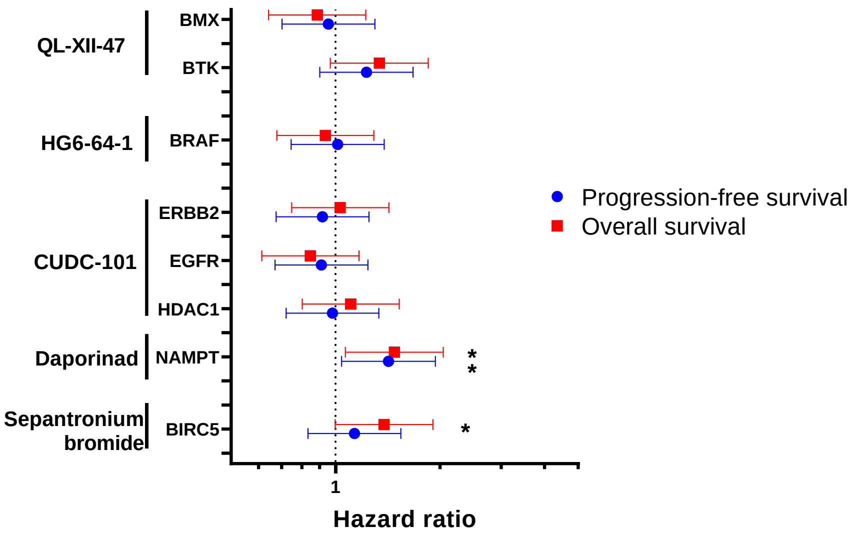

*Fig. S3: NAMPT correlations with EMT gene expression*

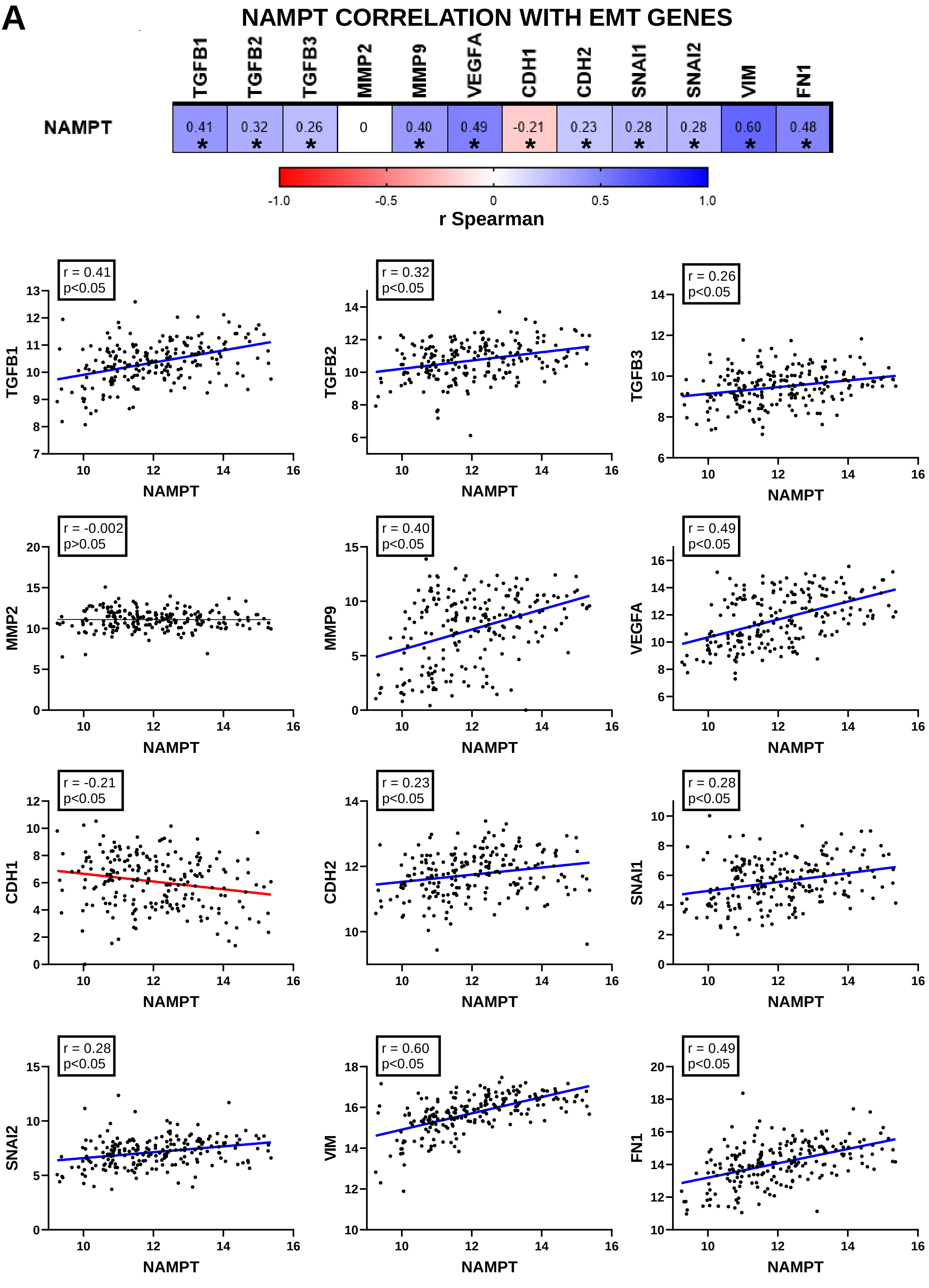

***Fig. S4: Predictive effect of Temozolomide according to the expression of targets for alternative drugs***

### TEMOZOLOMIDE VS. ALK PATHWAY

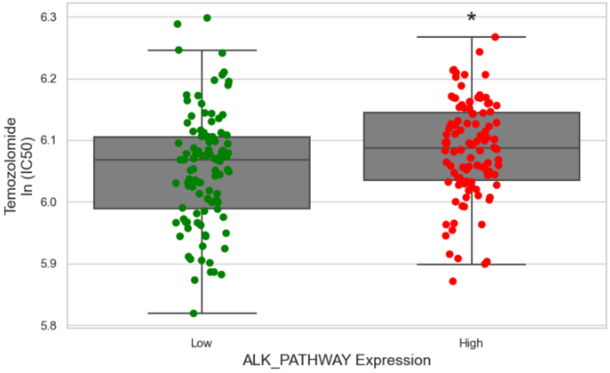


### TEMOZOLOMIDE VS. CTLA4_PATHWAY

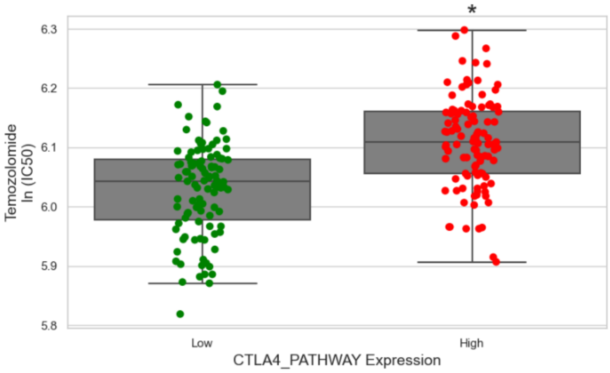


### TEMOZOLOMIDE VS. HDAC TARGETS

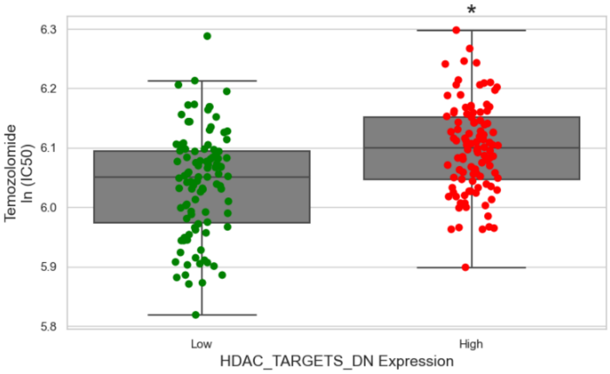


### TEMOZOLOMIDE VS. HER2 AMPLIFIED

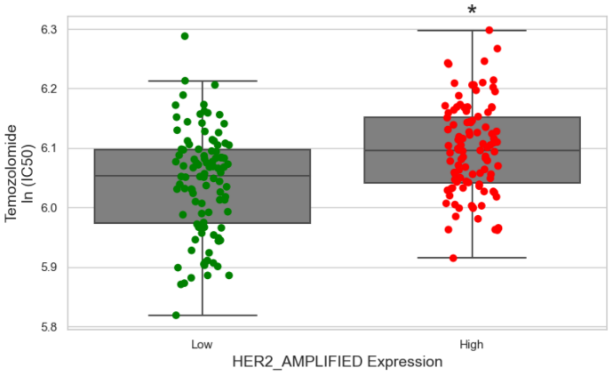


### TEMOZOLOMIDE VS. PI3K CASCADE

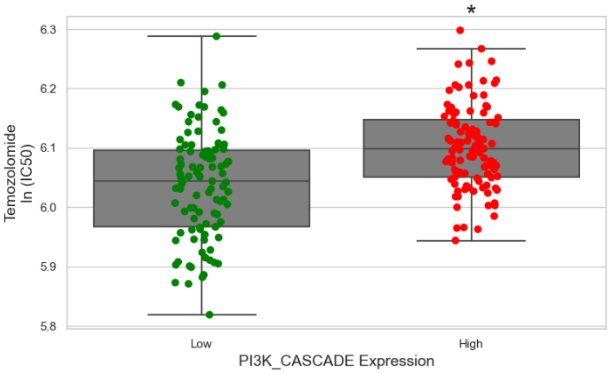


### TEMOZOLOMIDE VS. RB_PATHWAY

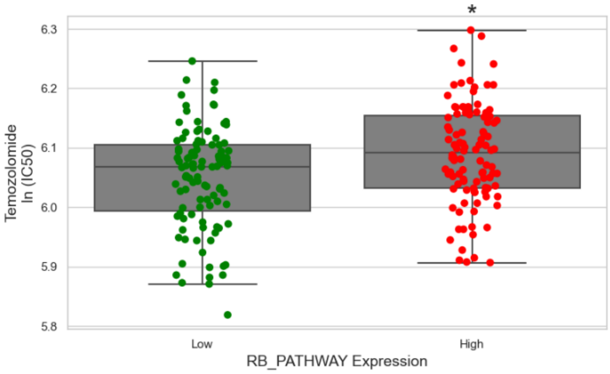


### TEMOZOLOMIDE VS. RETINOL METABOLISM

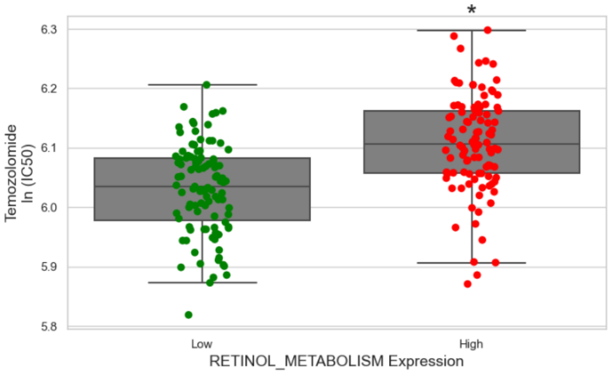

*Fig. S5: Correlations of Daporinad predictive effect with the expression of targets for alternative drugs*

### TEMOZOLOMIDE VS. ALK PATHWAY

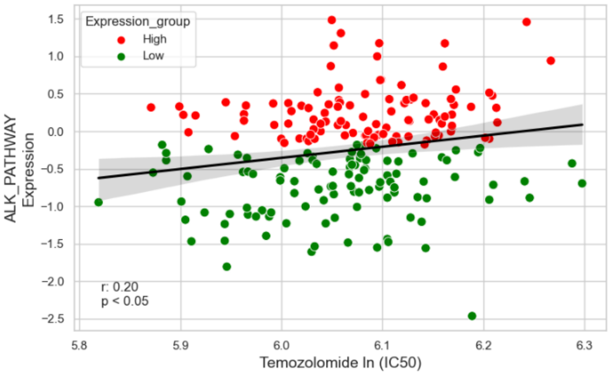


### TEMOZOLOMIDE VS. CTLA4_PATHWAY

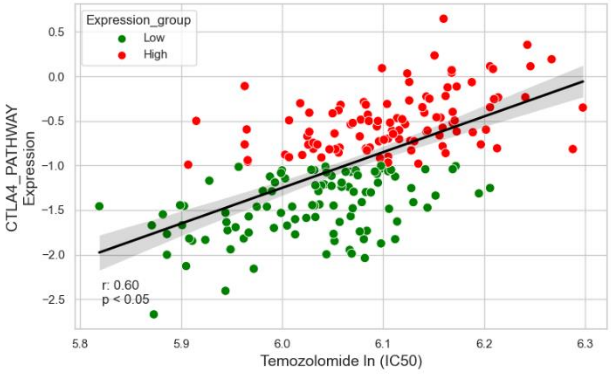


### TEMOZOLOMIDE VS. HDAC TARGETS

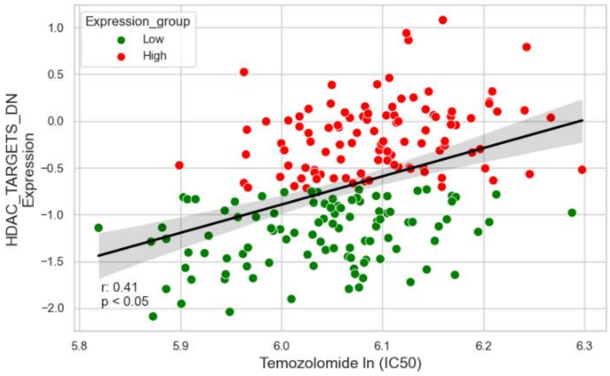


### TEMOZOLOMIDE VS. HER2 AMPLIFIED

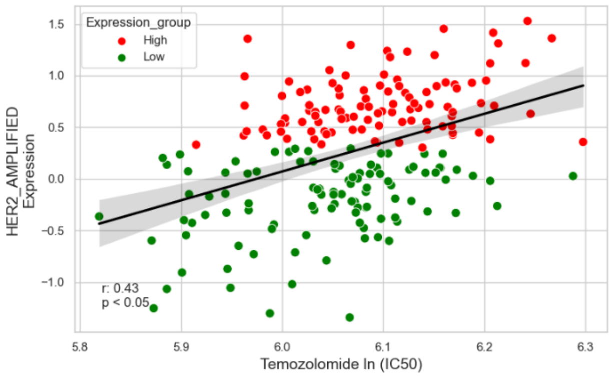


### TEMOZOLOMIDE VS. PI3K CASCADE

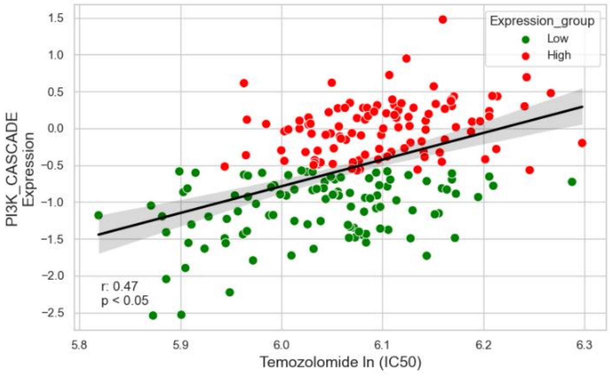


### TEMOZOLOMIDE VS. RB_PATHWAY

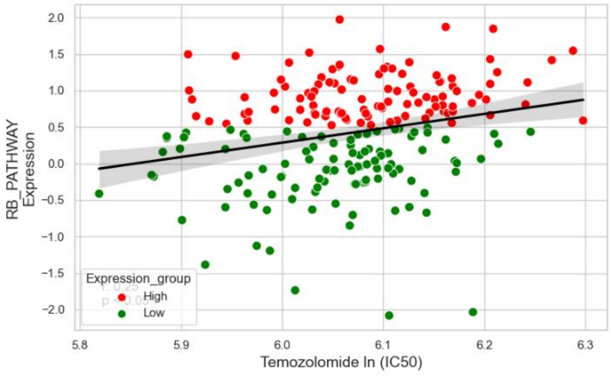


### TEMOZOLOMIDE VS. RETINOL METABOLISM

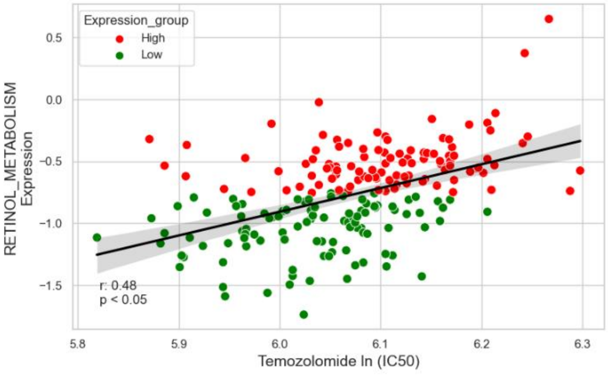

***Fig. S6: Predictive effect of Daporinad according to the expression of targets for alternative drugs***

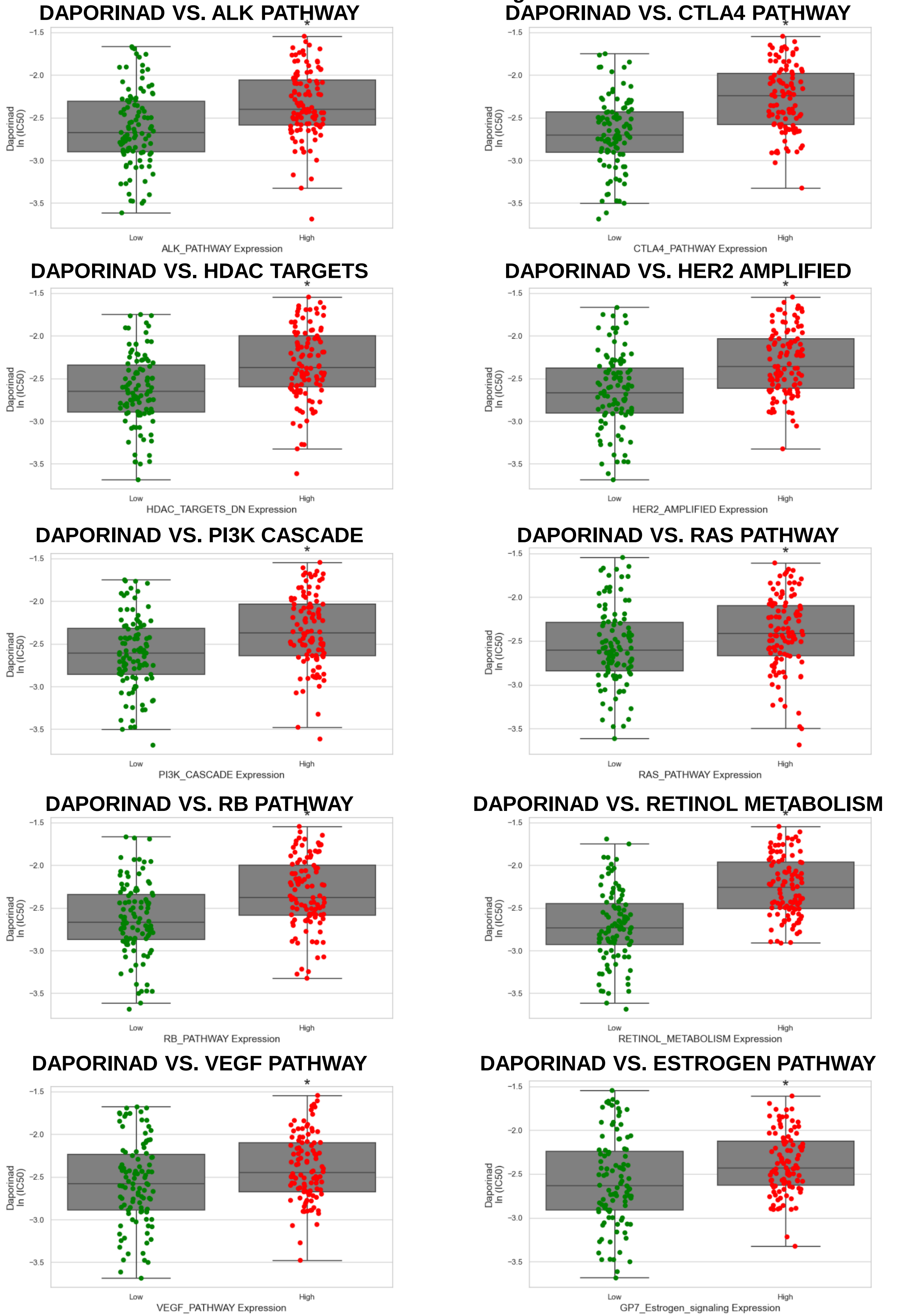

***Fig. S7: Correlations of Daporinad predictive effect with the expression of targets for alternative drugs***

### DAPORINAD VS. ALK PATHWAY

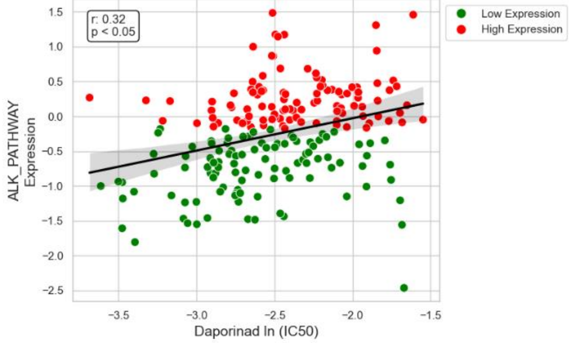


### DAPORINAD VS. CTLA4 PATHWAY

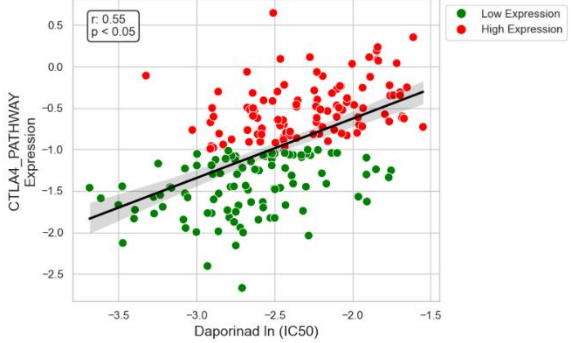


### DAPORINAD VS. HDAC TARGETS

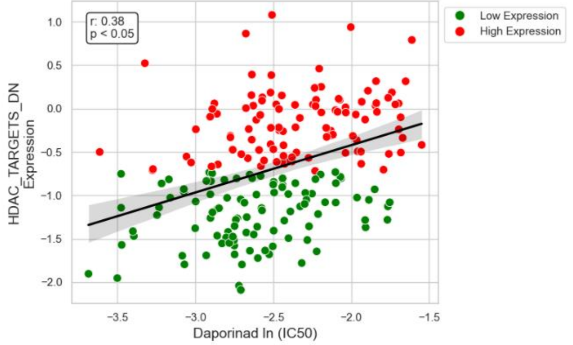


### DAPORINAD VS. HER2 AMPLIFIED

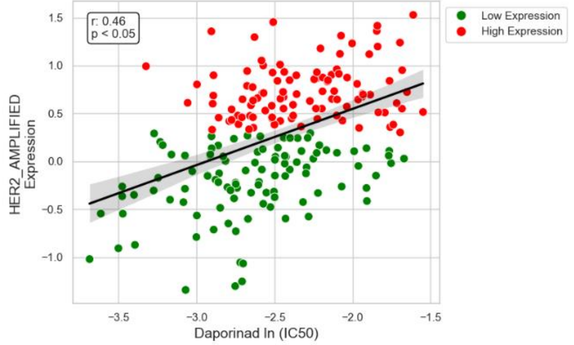


### DAPORINAD VS. PI3K CASCADE

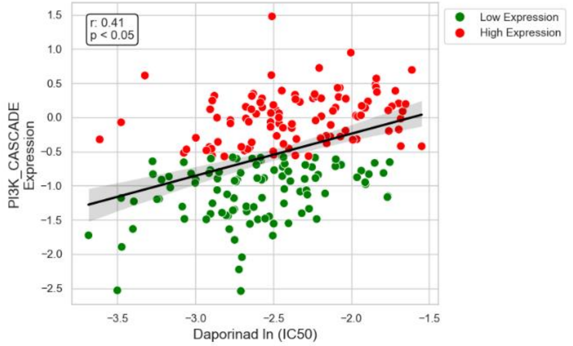


### DAPORINAD VS. RAS PATHWAY

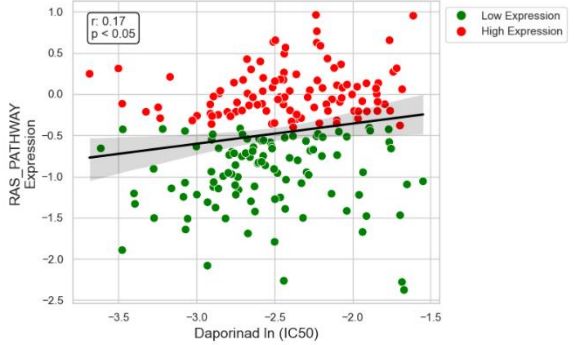


### DAPORINAD VS. RB PATHWAY

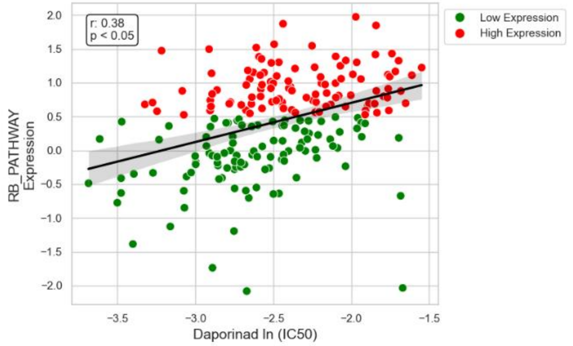


### DAPORINAD VS. RETINOL METABOLISM

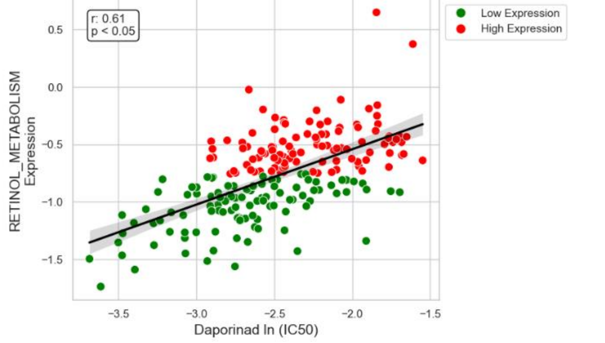


### DAPORINAD VS. VEGF PATHWAY

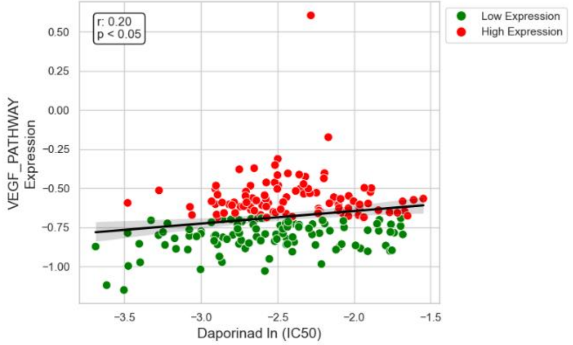


### DAPORINAD VS. ESTROGEN PATHWAY

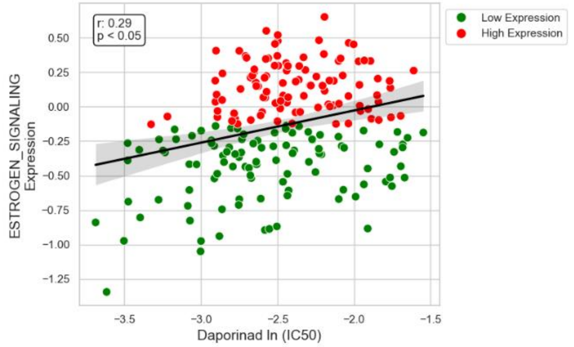
